# A glutamatergic DRN-VTA pathway for neuropathic pain and comorbid depression-like behavior modulation

**DOI:** 10.1101/2023.03.22.532931

**Authors:** Xin-Yue Wang, Wen-Bin Jia, Xiang Xu, Rui Chen, Liang-Biao Wang, Xiao-Jing Su, Peng-Fei Xu, Xiao-Qing Liu, Jie Wen, Yuan-Yuan Liu, Zhi Zhang, Xin-Feng Liu, Yan Zhang

## Abstract

Chronic pain causes both physical suffering and comorbid mental disorders such as depression. However, the neural circuits and molecular mechanisms underlying these maladaptive behaviors remain elusive. Here, we report a pathway from vesicular glutamate transporter3 neurons in the dorsal raphe nucleus to dopamine neurons in the ventral tegmental area (VGluT3^DRN^→DA^VTA^), of which population activity in response to innocuous mechanical stimuli and sucrose consumption, is respectively inhibited and attenuated by chronic neuropathic pain. Mechanistically, neuropathic pain dampens VGluT3^DRN^→DA^VTA^ glutamatergic transmission and DA^VTA^ neural excitability. VGluT3^DRN^→DA^VTA^ activation alleviates neuropathic pain and comorbid depression-like behavior (CDB) by releasing glutamate, which subsequently promotes DA release in the nucleus accumbens medial shell (NAcMed) and produces analgesia and antidepressant effects via D2 and D1 receptors, respectively. In addition, VGluT3^DRN^→DA^VTA^ inhibition produces pain-like hypersensitivity and depression-like behavior in healthy mice. These findings reveal a novel VGluT3^DRN^→DA^VTA^→D2/D1^NAcMed^ pathway in establishing and modulating chronic pain and comorbid depressive-like behavior.

## Introduction

Pain is a mixture of sensory/discriminative and motivational/affective experience. Nervous system elegantly interprets pain and pain relief as aversive and rewarding processes that trigger specific motivated behavioral responses. Acute pain results in adaptive escape/avoidance actions to prevent further tissue damage, and relief from acute pain is rewarding to facilitate learning of how to predict dangerous or rewarding situations in the future^1,2^. However, chronic pain, which affects near 20% of the people population word-wide, causes maladaptive physical suffering and mental disorders including the comorbid depression, which may conversely reinforce the intensity and duration of pain. The vicious cycle created by pain and depressive symptoms leads to a dilemma that medical management for chronic refractory pain is challenging. Understanding of the mechanisms underpinning chronic pain is essential for developing novel therapeutic strategies that lack adverse side effects.

The mesolimbic dopamine system, which mainly constitutes DA^VTA^ neurons projecting to the medial prefrontal cortex (mPFC) and nucleus accumbens (NAc), is perceived as a key machinery for encoding motivation, reward and aversion^3^. Dysfunction of dopamine system triggered by sustained pain states has been well investigated. Human and animal studies revealed that DA release was reduced in the NAc in both patients and rodents experiencing chronic pain^4–7^. Nonetheless, the circuit and molecular mechanisms underlying pain-induced adaptations within DA^VTA^ neurons and whether the adaptation of mesolimbic system is involved in comorbid depressive-like behavior during chronic pain are poorly studied.

Dopaminergic neurons in the VTA receive inputs from broad brain regions. Besides those that are known to be engaged in nocifensive information processing, including (e.g., the parabrachial nucleus (PBN), periaqueductal gray (PAG) and central amygdala (CeA)^8–11^), the dorsal raphe nucleus (DRN), one of the major sources of serotonin (5-HT) throughout the brain, also robustly projects to the VTA^12^. Apart from 5-HT, VTA-projecting DRN neurons also release co-transmitter glutamate which mediates reward^13,14^. Although ascending serotonergic neurons dysfunction has been implicated in depressive disorders including comorbid depressive symptoms during chronic pain^15^, whether maladaptation of glutamatergic transmission of DRN→VTA reward circuit underlies mesolimbic system dysfunction caused by neuropathic pain, and by extension underlies pain and CDB remains largely unknown.

In current study, using spared nerve injury (SNI) as a neuropathic pain model, we anatomically and functionally dissected a glutamatergic circuit from VGluT3^DRN^ to DA^VTA^ which undergoes maladaptive changes following neuropathic pain, resulting in reduced DA release in the NAcMed over NAc lateral shell (NAcLat). Activation of VGluT3^DRN^→DA^VTA^ pathway relieves pain and CDB via D2 receptor and D1 receptor in NAcMed, respectively. Conversely, VGluT3^DRN^→DA^VTA^ inhibition produces hypersensitivity and depressive-like behavior in healthy mice. Together, our results reveal the involvement of VGluT3^DRN^→DA^VTA^→NAcMed circuit adaptation in chronic pain and CDB.

## Results

### Neuropathic pain drives hypoexcitability of DA^VTA^ neurons

The mesolimbic dopamine system from the VTA is well known for reward and motivation processing^3,16^. Recent studies suggested that DA^VTA^ neurons also tuned to aversive stimuli including acute pain^17,18^. To assess whether dopamine dysfunction occurs during chronic pain, SNI was used as chronic neuropathic pain model in which long-lasting mechanical hypersensitivity develops after operation throughout the experimental timeline of 6 weeks (6W) (Supplementary Fig. 1a, b). In line with a previous study^15^, SNI mice developed CDB, which was assessed by the sucrose preference test (SPT), a standard assay for assessing anhedonia which is a key defining feature of depression^19^, at relatively late-stage (SNI 6W), rather than at early-stage of neuropathic pain (SNI 2W) (Supplementary Fig. 1c).

To investigate changes in the dynamic activity of DA^VTA^ neurons associated with nerve injury-induced pain-like hypersensitivity and CDB, *in vivo* fiber photometry synchronized with a video-tracking system was performed to monitor Ca^2+^ fluctuations upon hindpaw von Frey stimulation and sucrose consumption, respectively. First, we injected Cre-dependent adeno-associated virus (AAVs) expressing the fluorescent Ca^2+^ indicator GCaMP6m into the VTA of *DAT-Cre* mice (Fig. 1a, b). *Post hoc* staining showed that 93.7% of GCaMP6m-expressing neurons were Tyrosine hydroxylase (TH)-positive, indicative of high specificity of viral targeting (Fig. 1c). To characterize the activity of DA^VTA^ neurons associated with mechanical hypersensitivity, innocuous punctate mechanical stimuli (0.4 g von Frey filament), which rarely evokes paw withdrawal responses in naïve mice but causes frequent withdrawal and aversion after SNI, was applied to the hind paw (Fig. 1d and Supplementary Fig. 2a, b). Interestingly, we observed a significant DA^VTA^ neural activation which was time-locked to von Frey stimulation in pre-SNI healthy mice, whereas the same stimuli evoked an overall inhibition of calcium transients in post-SNI 2W mice (Fig. 1e). We next examined population activity of DA^VTA^ neurons associated with CDB. In pre-SNI mice, sucrose consumption induced the dramatic Ca^2+^ elevation, indicating a prominent role of DA^VTA^ neurons in reward and motivation processing (Fig. 1f, g) as previously reported^20^. In consistent with our behavioral measurements (Supplementary Fig. 1c), such activity increment of DA^VTA^ neurons was maintained at 2 weeks post SNI (Supplementary Fig. 3a, b). However, in post-SNI 6W mice which developed the CDB, the activation of DA^VTA^ neurons upon sucrose licking was significantly decreased (Fig. 1f, g). These results established a potential link between dysfunctional adaptation in the DA^VTA^ neural activity and nerve injury-induced pain-like hypersensitivity and CDB.

**Fig 1.**
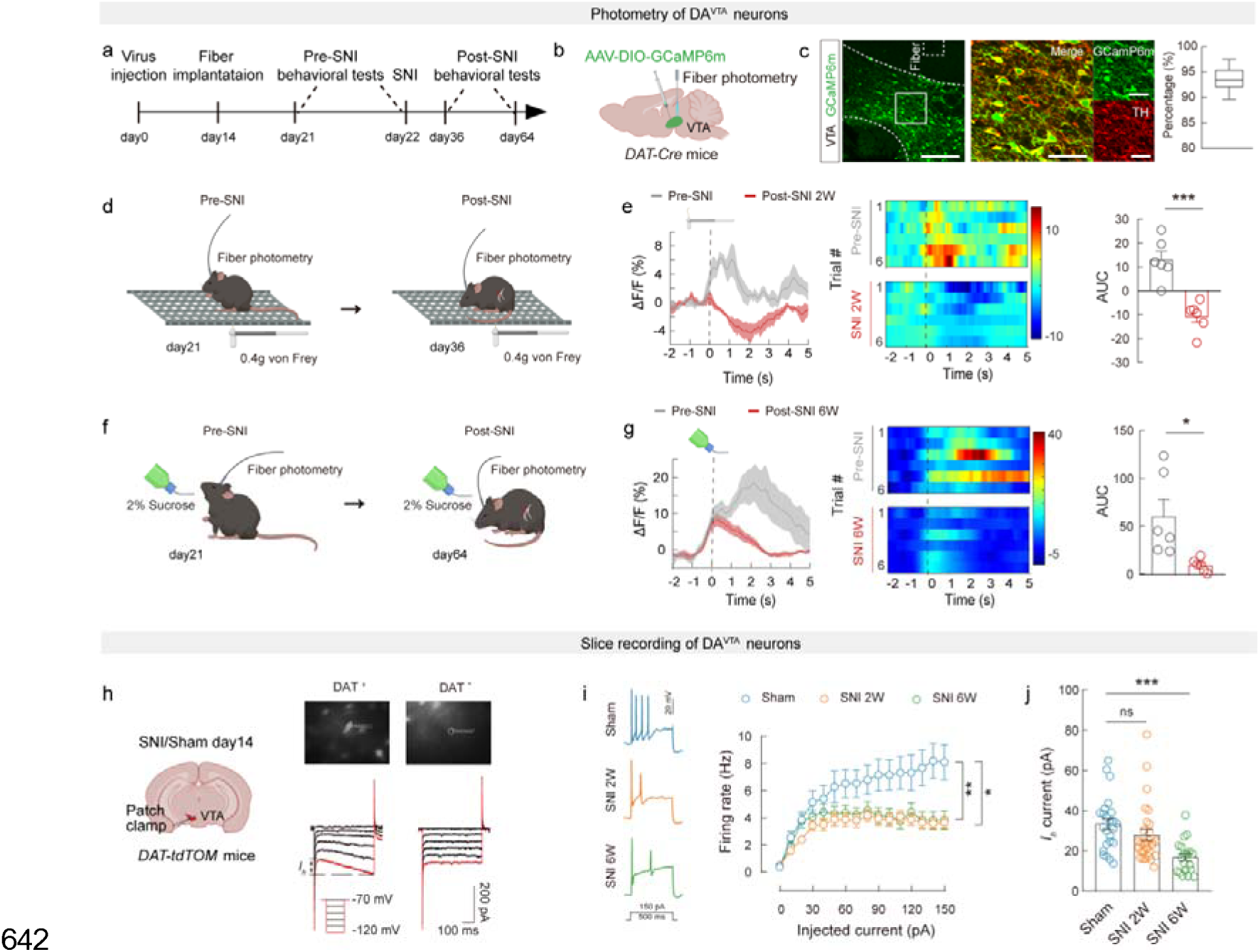
Dysfunctional adaptation in the DA^VTA^ neural activity in mice with chronic pain. **a,** Schematic of the experimental design. **b**, Schematic diagram of viral injection and fiber photometry of DA^VTA^ neurons. **c**, Representative images of viral expression and track of the optical fiber in the VTA (left) in *DAT-Cre* mice. Summary data for percentage of GCaMP6m-expressed neurons co-localized with TH immunofluorescence within the VTA, n = 9 sections from three mice (right). Scale bars, 200 μm (left) and 50 μm (right). **d**, **f**, Schematic of the experimental design for von Frey (d) and sucrose test (f). **e**, **g**, Averaged responses (left), heatmaps (middle) and area under the curve (AUC) during 0-5 s (right) showing Ca^2+^ responses evoked by von Frey stimulation in SNI 2W mice compared with pre-SNI mice (**e**) and sucrose licking in SNI 6W mice compared with pre-SNI mice (**g**). **h**, Schematic of the electrophysiological recordings in acute slices (left) and typical currents induced by hyperpolarizing voltage steps recorded from the DAT^+^ (tdTOM positive) and DAT^-^ (tdTOM negative) neurons (right). **i**, Representative traces (left) and statistical data (right) of firing rate recorded from Sham, SNI 2W and SNI 6W mice. **j**, Statistical data of hyperpolarization-activated currents (*Ih*) at-120 mV recorded from DAT^+^ neurons. Significance was assessed by two-tailed paired Student’s *t*-test in (**e, g**), two-way ANOVA followed by Bonferroni’s multiple comparisons test between groups in (**i**), and one-way ANOVA followed by Bonferroni’s multiple comparisons test in (**j**). For (**c**), data are shown as box and whisker plots (medians, quartiles (boxes) and ranges minimum to maximum (whiskers); For (**d-j**) data are presented as the mean ± s.e.m. \**P* < 0.05, \*\**P* < 0.01, \*\*\**P* < 0.001, not significant (ns). Details of the statistical analyses are presented in Supplementary Table 1.

Upon observing changes in the population activity of VTA^DA^ neurons during neuropathic pain, we went further to analyze the underlying physiological mechanisms at single neuron level by whole-cell patch clamp recordings in brain slices. To visualize dopaminergic neurons, transgenic *DAT-tdTOM* mice was generated by crossing *DAT-Cre* mice with *Ai14* strain (Fig. 1k). DAT^+^ neurons from the VTA displayed a typical hyperpolarization-activated cation current (*Ih*) (Fig. 1h) and a lower excitability profile to depolarizing current steps when compared to DAT^-^ neurons (Fig. 1h and Supplementary Fig. 4a) which are likely to be GABAergic^21,22^. As expected, DAT^+^ neurons in slices from SNI 2W mice showed significant decrease in the firing rate following 500-ms current injection with increments of 10 pA from 0 to 150 pA compared to sham controls (Fig. 1i). In contrast, the excitability of DAT^-^ neurons was comparable between two groups (Supplementary Fig. 4a, b). We then postulated that further reduction of firing rate of DA^VTA^ neurons in SNI 6W mice caused CDB manifestation. Unexpectedly, the DA^VTA^ neurons from SNI 6W mice displayed similar firing rate as those from SNI 2W mice (Fig. 1i), implying other contributors to the depression-like behavior. Given the unique role of *Ih* in modulating intrinsic excitability and temporal integration of synaptic input^23^, we analyzed *Ih*, and, intriguingly, observed dramatic *Ih* reduction in DAT^+^ neurons from SNI 6W mice in comparison with sham controls and SNI 2W mice (Fig. 1j).

From above, population activity and intrinsic excitability of DA^VTA^ neurons are blunted during chronic neuropathic pain. We thus examined whether artificial activation of the DA^VTA^ neurons by chemogenetic strategy would be sufficient to ameliorate neuropathic pain and CDB. For this, *DAT-Cre* mice were injected with a Cre-dependent excitatory DREADD (AAV-DIO-hM3Dq-mCherry), or a control vector (AAV-DIO-mCherry) into the ipsi-lateral VTA (Supplementary Fig. 5a, b). High overlapping of hM3Dq-mCherry-expressing neurons with TH staining proved high viral efficacy and specificity (Supplementary Fig. 5b). Two weeks after nerve injury, chemogenetic activation of the DA^VTA^ neurons was sufficient to rescue the mechanical and thermal hypersensitivity in CNO-treated hM3Dq mice (Supplementary Fig. 5c, d), whereas it had no effect in control (saline-treated hM3Dq mice and mCherry mice) animals. This is consistent with a previous study^24^. Moreover, 6 weeks after SNI, chemogenetic activation of the DA^VTA^ neurons remarkably alleviated SNI-induced CDB (Supplementary Fig. 5e).

## Neurochemical and electrophysiological characterization of the DRN→VTA circuit

To investigate the potential neural circuit mechanism underlying the neuropathic pain-induced hypoexcitability of DA^VTA^ neurons, rabies virus (RV)-based retrograde trans-monosynaptic tracing method was conducted by injecting Cre-dependent helper viruses (AAV-EF1α-DIO-TVA-GFP and AAV-EF1α-DIO-RVG) into the VTA of *DAT-Cre* or *GAD2-Cre* mice (Fig. 2a). Two weeks later, EnvA-pseudotyped RV-ΔG-DsRed was injected into the same site. In *DAT-Cre* mice, substantial DsRed-labeled neurons were detected in the DRN, laterodorsal tegmentum (LDTg), PBN, lateral habenular nucleus (LHb) and CeA (Fig. 2b and Extended Data Fig. 6b). Because glutamatergic projections from the DRN to VTA strongly mediate reward^13,14,25,26^, we thus particularly focused on this circuit in chronic neuropathic pain and comorbid depression. Immunofluorescence staining showed that ∼62% of DsRed-labeled neurons in the DRN co-localized with VGluT3 (Fig. 2d), a vesicular transporter which concentrates glutamate into synaptic vesicles^27^ and is essential for glutamate release to signal reward^14,25^. In comparison, less than 30% of DsRed-labeled neurons co-expressed VGluT3 in the DRN of *GAD2-Cre* mice (Fig. 2c, d, and Supplementary Fig. 6c, d). Anterograde trans-monosynaptic tracing strategy by injecting AAV2/1-Cre into the DRN and AAV-DIO-mCherry into the VTA of *C57* mice further verified the anatomical connection of DRN→VTA circuit (Supplementary Fig. 7a). Moreover, ∼71% of DRN-targeted VTA neurons labeled by mCherry co-stained with TH antibody, and ∼35% of them expressed GABA (Supplementary Fig. 7b, c). These morphological data demonstrate the presence of VGluT3^DRN^→DA^VTA^ circuit, and the VGluT3^DRN^ neurons send major projections to DA^VTA^ neurons instead of GABA^VTA^ neurons.

**Fig 2.**
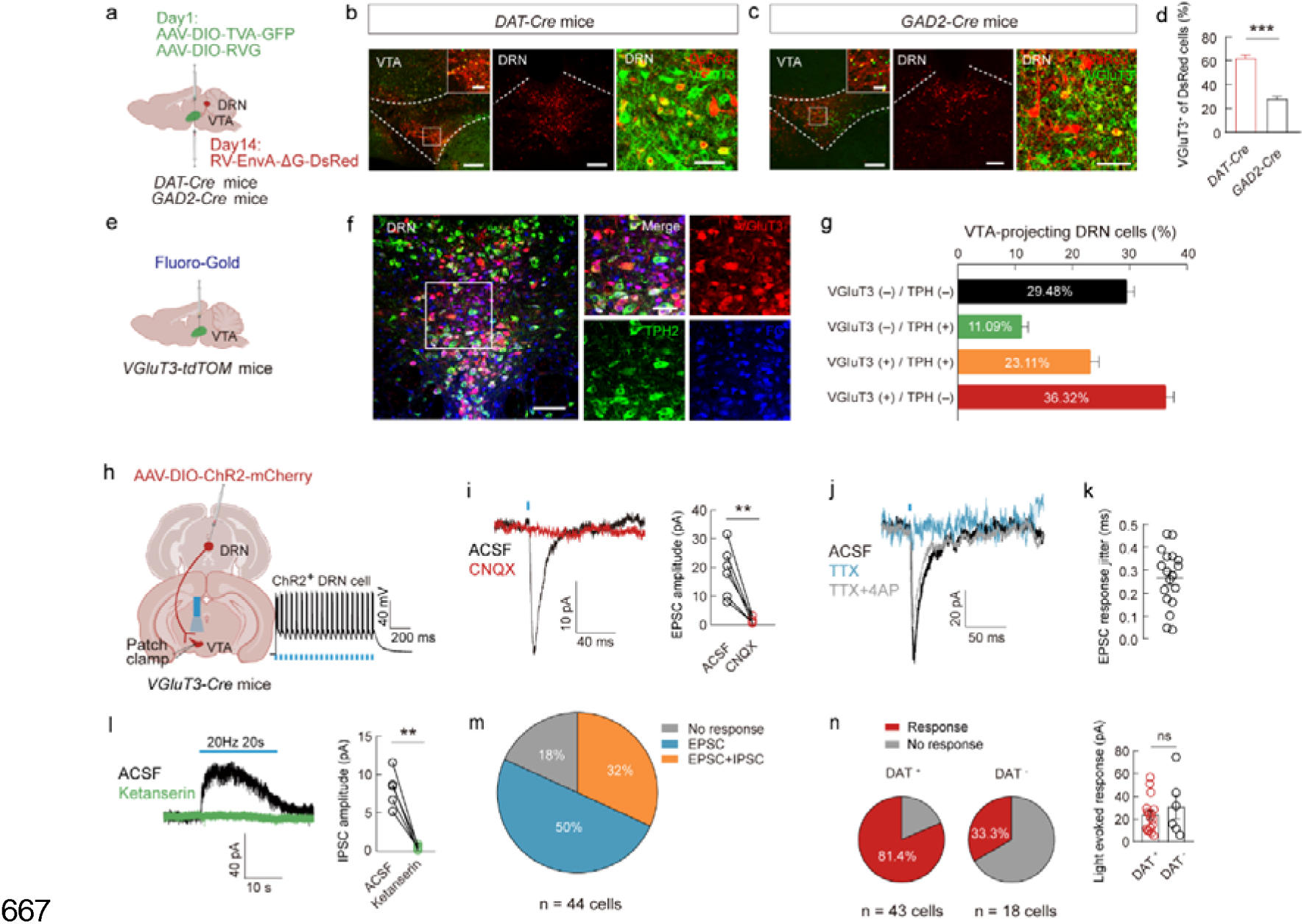
Defining of VGluT3^DRN^→DAT^VTA^ circuits. **a**, Schematic of the Cre-dependent retrograde monosynaptic tracing strategy in *DAT-Cre* and *GAD2-Cre* mice. **b**, **c**, Representative images of the starter cells in the VTA (left) and RV-DsRed-labeled cells in the DRN (middle) which co-localize with VGluT3 immunofluorescence (right) from *DAT-Cre* (b) mice and *GAD2-Cre* mice (c). Starter cells (yellow) co-express AAV-DIO-TVA-GFP, AAV-DIO-RVG (green) and Δ m (left, middle) and 50 μm (right). **d**, Summary data for percentage of DsRed-labeled neurons that expressed VGluT3 of *DAT-Cre* and *GAD2-Cre* mice, n = 9 sections from three mice. **e**, Schematic diagram of retrograde tracer FG injected into the VTA. **f**, Typical images of the FG (bule) neurons co-localize with TPH2 immunofluorescence (green) in the DRN of *VGluT3-tdTOM* mice. Scale bars, 100 μm (left) and 50 μm (right). **g**, Percentage of FG-labeled neurons that expressed VGluT3 and TPH2 in the DRN, n = 15 sections from four mice. **h**, Schematic showing viral injection and VTA electrophysiological recordings in acute slices from *VGluT3-Cre* mice (left). Viral expression efficacy proved by light (473 nm, 20 Hz)-evoked action potentials of ChR2-mCherry-expressing neuron (right). **i**, Representative traces and summary data for light-evoked EPSCs of VTA neurons before (ACSF) and after CNQX (10 μ treatment. **j**, Representative traces of light-evoked EPSCs of the VTA neurons before (ACSF) and after TTX (1 μM) or TTX and 4-AP (100 μM) treatment. **k**, Summary data for jitter of light-evoked EPSCs recorded from VTA neurons. **l**, Representative traces and summary data for light-evoked (473 nm, 20 s, 20 Hz) slow IPSCs of VTA neurons before and after ketanserin (10 μM) treatment. **m**, Pie chart showing the distribution of light-evoked response types in DRN-targeted VTA neurons. **n**, Summary data for percentage (left) and amplitudes of light-evoked fast EPSCs recorded from DAT^+^ and DAT^-^ neurons. Significance was assessed by two-tailed unpaired Student’s *t*-test in (**d**, **n**), two-tailed paired Student’s *t*-test in (**i**, **l)**. All data are presented as the mean ± s.e.m. \*\**P* < 0.01, \*\*\**P* < 0.001, not significant (ns). Details of the statistical analyses are presented in SupplementaryTable 1.

We next systematically analyzed the neurochemical identity of VTA-projecting VGluT3^DRN^ (VGluT3^DRN→VTA^) neurons by injecting Fluoro-Gold (FG) into the VTA of *VGluT3-tdTOM* mice (Fig. 2e). At 7 days after injection, we detected FG-labeled neurons in the DRN, and observed that ∼60% of FG^+^ neurons contained VGluT3 (∼36% with VGluT3 only and ∼23% with both 5-HT and VGluT3) (Fig. 2f, g). In comparison, only 11% of FG-labeled neurons expressed 5-HT alone. Thus, the majority of VTA-projecting DRN neurons are VGluT3-expressing, some of which co-express 5-HT.

The functional connection of the VGluT3^DRN^→VTA circuit was further characterized by electrophysiology. We injected Cre-dependent AAV-DIO-channelrhodopsin 2 (ChR2)-mCherry into the DRN of *VGluT3-Cre* mice to enable selective activation of the VGluT3^DRN→VTA^ terminals to examine postsynaptic responses in the VTA (Fig. 2h). Current clamp recordings on ChR2-expressing DRN neurons showed that 20 Hz blue light stimulation reliably evoked action potentials without failure, suggesting the functional viral transduction (Fig. 2h). When recordings were performed on VTA neurons, excitation of the VGluT3 afferents by single-pulse light produced frequently (22 of 44 neurons, 50%) fast-onset excitatory postsynaptic currents (fast EPSCs) (Fig. 2i, m), but rare (1 of 44 neurons, data not shown), if any, fast-onset inhibitory postsynaptic currents (fast IPSCs). These EPSCs were almost fully eliminated by bath application of CNQX (10 μM), a selective AMPA/kainate receptor antagonist (Fig. 2i). Moreover, EPSCs could be blocked by tetrodotoxin (TTX) and restored by bath application of the potassium channel blocker 4-aminopyridine (4-AP) (Fig. 2j). Low-response jitter was also observed in light-evoked EPSCs (Fig. 2k). Taken together, these data suggest that VTA neurons receive monosynaptic glutamatergic inputs from VGluT3^DRN^ neurons. When we stimulated the VGluT3^DRN→VTA^ afferents by prolonged blue light (20 Hz for 20 s), ∼32% (14 of 44) of VTA neurons displayed slow outward currents (slow IPSCs) (Fig. 2l, m). The slow IPSCs can be abolished by ketanserin, a selective antagonist for 5-HT2A and 5-HT2c receptors (Fig. 2l). Of note, the 32% of VTA neurons with slow IPSCs concurrently had detectable fast EPSCs, which resembles above neurochemical study. Altogether, VTA neurons receive both glutamatergic and serotonergic inputs from VGluT3^DRN^ neurons. Furthermore, we specifically recorded DA^VTA^ neurons, which were validated by the presence of a hyperpolarization-activated cation current (*Ih)*, and found that ∼81% of them displayed fast EPSCs following VGluT3^DRN→VTA^ afferents excitation, whereas only ∼33% of *Ih*-negative neurons with fast EPSCs were observed (Fig. 2n). These findings together with the morphological data prove that VGluT3^DRN^ neurons make functional connection preferentially with DA^VTA^ neurons.

## Neuropathic pain dampens VGluT3^DRN^→DA^VTA^ circuit activity

To investigate a potential association of the DRN→VTA pathway adaptation with nerve injury-induced pain-like hypersensitivity and CDB, we used fiber photometry to measure calcium dynamics from VGluT3^DRN→VTA^ terminals in response to hind paw mechanical stimuli and sucrose consumption. To this end, *VGluT3-Cre* mice received DRN injection with AAV-DIO-GCaMP6m and VTA implantation with optical fiber (Fig. 3a, b). Under physiological condition, we observed an elevation in calcium activity upon the delivery of 0.4 g von Frey filament, whereas the same stimuli induced large inhibition in calcium activity in post-SNI 2W mice (Fig. 3d). We further monitored the calcium signal of VGluT3^DRN→VTA^ terminals in response to sucrose consumption. In healthy pre-SNI mice, sucrose licking significantly increased calcium transients of VGluT3^DRN→VTA^ terminals. Two weeks after SNI, sucrose consumption increased calcium transients to the same extent as pre-SNI mice (Supplementary Fig. 8a, b). However, in post-SNI 6W mice that displayed CDB, sucrose licking triggered calcium signal at a much lower level (Fig. 3e). These findings demonstrate that VGluT3^DRN→VTA^ afferents activity is altered by SNI.

**Fig 3.**
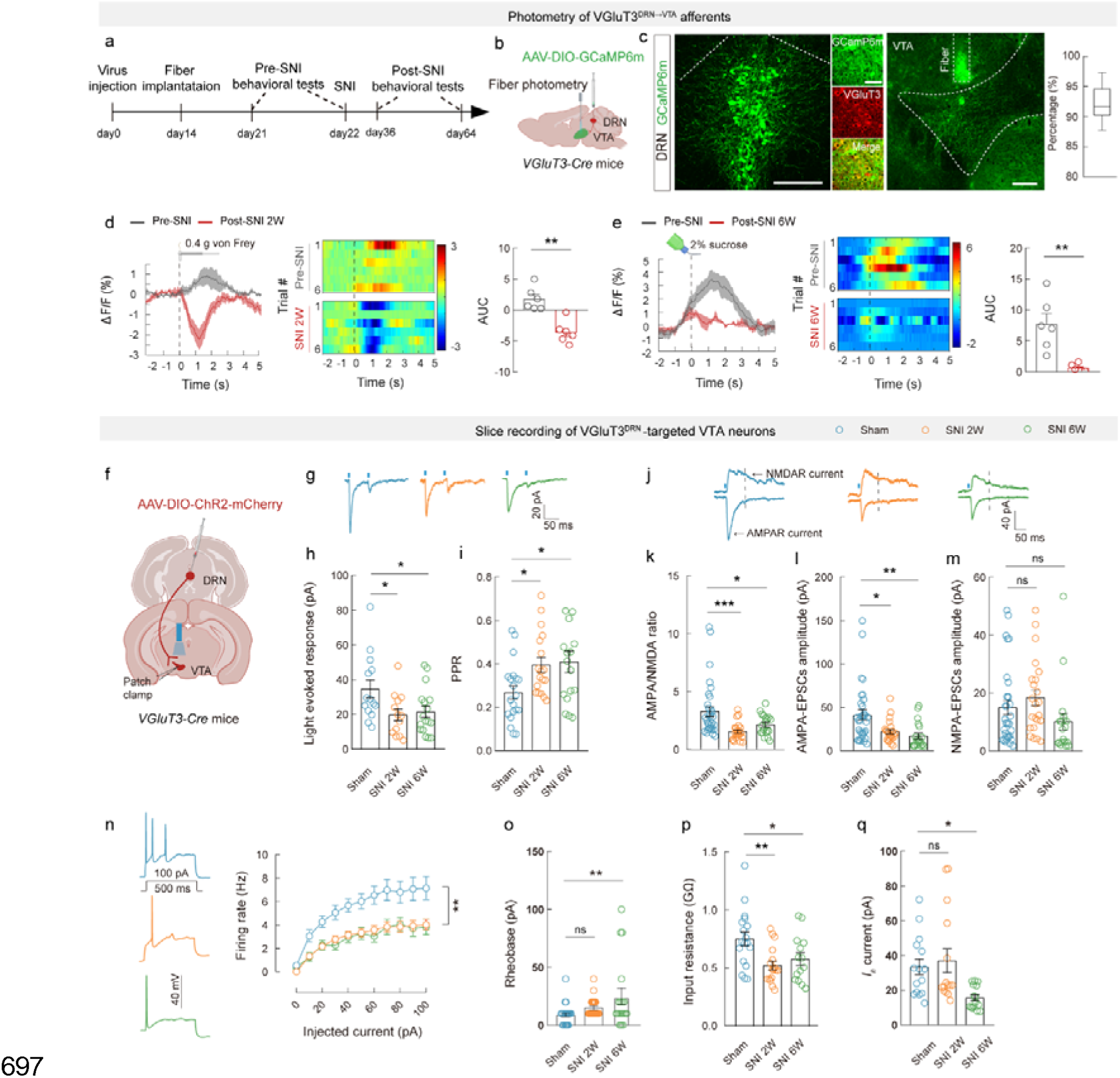
Dampened activity of VGluT3^DRN^→DA^VTA^ circuit in mice with chronic pain. **a**, Schematic of the experimental design. **b**, Schematic diagram of viral injection and fiber photometry of VGluT3^DRN→VTA^ terminals in *VGluT3-Cre* mice. **c**, Representative images of viral expression in the DRN and track of the optical fiber in the VTA (left). Summary data for percentage of GCaMP6m-expressed neurons co-localized with VGluT3 immunofluorescence within the DRN, n = 9 sections from three mice (right). Scale bars, 200 μm (left, right) and 50 μm (middle). **d**, **e**, Averaged responses (left), heatmaps (middle) and AUC during 0-5 s (right) showing Ca^2+^ responses evoked by von Frey stimulation in SNI 2W mice compared with pre-SNI mice (d) and sucrose licking in SNI 6W mice compared with pre-SNI mice (e). **f**, Schematic showing viral injection and the electrophysiological recordings in acute slices from *VGluT3-Cre* mice. **g**, Representative traces of paired-pulse ratio (PPR) at DRN-VTA pathway recorded from DA^VTA^ neuronsof Sham, SNI 2W, and SNI 6W mice. **h**, **i**, Summary data for the amplitudes (h) and PPR of light-evoked currents (i) recorded from the VGluT3^DRN^-targeted DA^VTA^ neurons. **j**, Representative traces of AMPA-EPSCs and NMDA-EPSCs evoked by optogenetic activation of DRN inputs at-70 and +40 mV, respectively. Time points for determination of AMPA and NMDA receptor currents were indicated with dashed lines. **k**-**m**, Statistics of AMPA/NMDA ratio (left), AMPA-EPSCs amplitude (middle) and NMDA-EPSCs amplitude (right) at DRN-VTA pathway. **n**-**q**, Representative traces and statistical data for action potential firing (**n**), rheobase (**o**), input resistance (**p**) and *Ih* at-120 mV (**q**) recorded from VGluT3^DRN^-targeted postsynaptic DA^VTA^ neurons. Significance was assessed by two-tailed paired Student’s *t*-test in (**d**, **e**), one-way ANOVA followed by Bonferroni’s multiple comparisons test in (**h**, **i**, **k-m**, **o-q**), and two-way ANOVA followed by Bonferroni’s multiple comparisons test in (**n**). For (**c**), data are shown as box and whisker plots (medians, quartiles (boxes) and ranges minimum to maximum (whiskers); For (**d-q**) data are presented as the mean ± s.e.m. \**P* < 0.05, \*\**P* < 0.01, \*\*\**P* < 0.001, not significant (ns). Details of the statistical analyses are presented in Supplementary Table 1.

To further explore the plastic changes in VGluT3^DRN→^DA^VTA^ circuit activity during chronic neuropathic pain, we performed patch-clamp recordings from VTA neurons. Nerve injury did not affect the proportion of VGluT3^DRN→VTA^ terminals activation-evoked postsynaptic responses in both post-SNI 2W and post-SNI 6W mice versus naïve mice (Supplementary Fig. 9a, b). Notably, we did not observe any difference in amplitudes of slow IPSCs between naïve mice and post-SNI mice (Supplementary Fig. 9c, d), whereas we observed a significant attenuation in amplitudes of fast EPSCs recorded from *Ih*-positive DA^VTA^ neurons in SNI mice compared to sham controls (Fig. 3f, h). The compromised glutamatergic transmission may arise from either decreased presynaptic glutamate release from VGluT3^DRN→VTA^ neurons or/and undermined postsynaptic response to glutamate of VTA neurons. We firstly assessed paired-pulse ratio (PPR) of light-evoked fast EPSCs, which was well known to be inversely correlated with the presynaptic transmitter release. We found that PPR (ISI 50 ms) of DA^VTA^ neurons was significantly increased in SNI mice compared to sham controls (Fig. 3g, i), implying attenuated presynaptic glutamate release in VGluT3^DRN→VTA^ neurons. Next, we investigated the ratio of AMPA receptor-mediated EPSCs (AMPA-EPSCs) over NMDA-EPSCs, an indicator of postsynaptic plasticity, and detected a smaller ratio in SNI mice which is likely caused by selective attenuation of AMPA-EPSCs rather than NMDA-EPSCs (Fig. 3j-m).

We also analyzed the intrinsic excitability of VGluT3^DRN^-targeted postsynaptic DA^VTA^ neurons which were identified by the presence of *Ih* currents and light-evoked fast EPSCs. The input-output plot showed lowered firing rate of DA^VTA^ neurons in slices from both SNI 2W and SNI 6W mice compared to sham controls (Fig. 3n). Correspondingly, the rheobase, calculated by minimal current required to evoke an action potential, was increased (Fig. 3o). Consistent with the decreased excitability, a significant decrease in the input resistance was also observed in both SNI 2W and SNI 6W mice (Fig. 3p), whereas significant reduction in *Ih* was only detected in SNI 6W mice with CDB (Fig. 3q). In summary, these data show that SNI dampens synaptic strength of the VGluT3^DRN^ DA^VTA^ pathway via both presynaptic and postsynaptic mechanisms.

## The VGluT3^DRN→VTA^ terminals modulate chronic pain and comorbid depression

Considering that synaptic transmission of VGluT3^DRN^→DA^VTA^ circuit was compromised during chronic neuropathic pain, we postulated that artificial activation of the VGluT3^DRN→VTA^ projection would alleviate pain-like hypersensitivity and CDB. We injected *VGluT3-Cre* mice with AAV-DIO-ChR2-mCherry or AAV-DIO-mCherry into the DRN and implanted optical fibers above the VTA (Fig. 4a, b). We found that optogenetic activation of VGluT3^DRN→VTA^ terminals significantly elevated paw withdrawal threshold in the von Frey test and paw withdraw latency in the Hargreaves test of SNI 2W mice (Fig. 4c, d), whereas exerted no effect in sham control mice. The locomotion assessed by open-field test was unaffected by VGluT3^DRN→VTA^ terminals activation (Supplementary Fig. 10a, b). These results indicate that VGluT3^DRN→VTA^ terminals excitation alleviated neuropathic pain, without affecting basal mechanical and thermal nociception.

**Fig 4.**
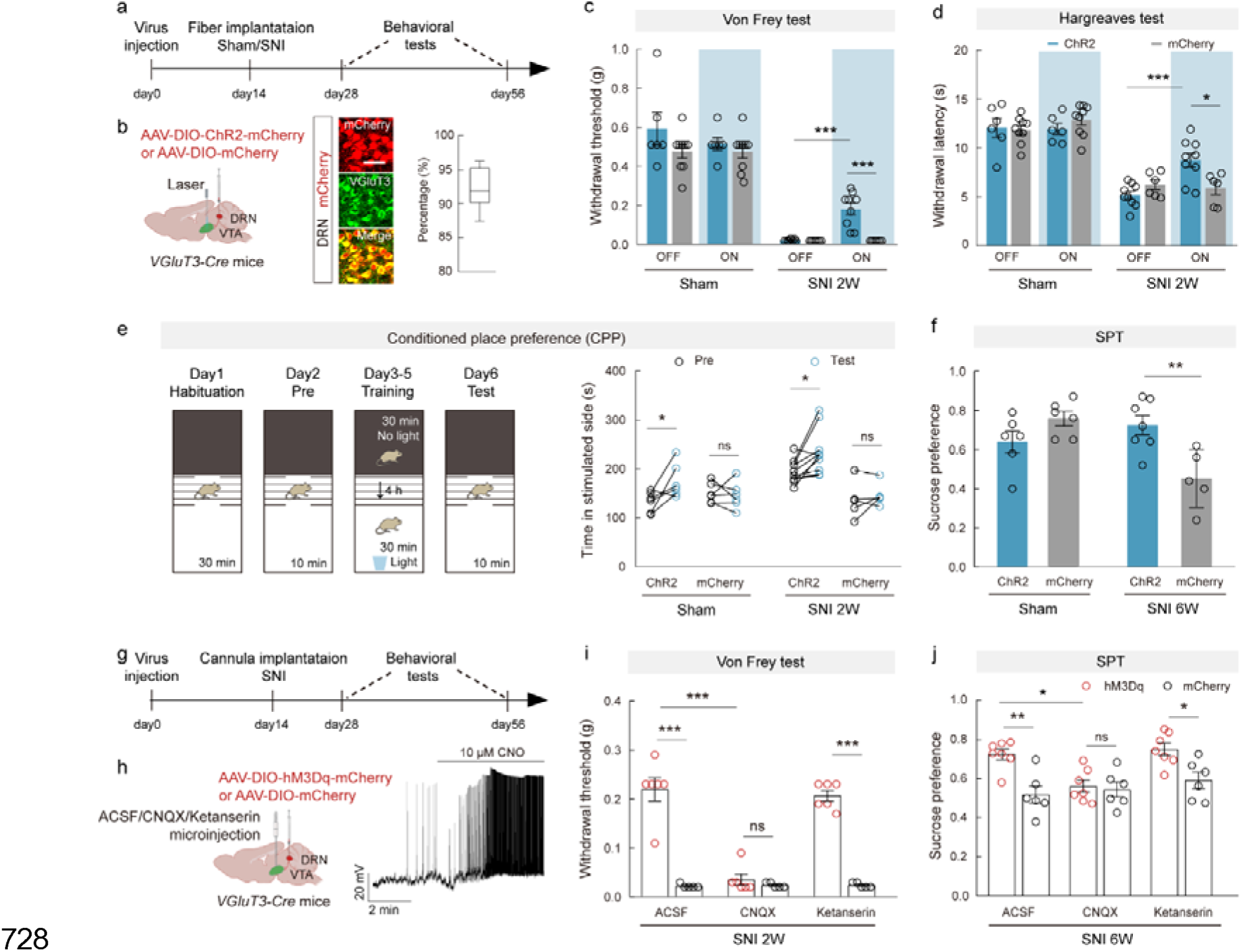
Glutamate is essential for analgesic and anti-depressive effects by the VGluT3^DRN→VTA^ neural activation. **a, g**, Schematic of the experimental design. **b**, Schematic of DRN injection of AAV-DIO-ChR2-mCherry/AAV-DIO-mCherry and VTA optical fiber implantation in *VGluT3*-Cre mice (left). Representative images (middle) and summary data (right) for percentage of mCherry-expressed neurons co-localized with VGluT3 immunofluorescence in the DRN, n = 9 sections from three mice. Scale bar, 50 μm. **c**, **d**, Mechanical paw withdrawal threshold (left) and thermal paw withdrawal latency (right) of ChR2-mCherry-expressing and mCherry-expressing Sham or SNI mice with (on) or without (off) optogenetic activation of VGluT3^DRN→VTA^ terminals. **e**, Experimental design of conditional place preference (CPP) test (left) and quantification of CPP before (Pre) and after (Test) training of the Sham or SNI mice injected with AAV-DIO-ChR2-mCherry or control AAV-DIO-mCherry (right). **f**, Preference for sucrose in the SPT. **h**, Schematic diagram of DRN injection of AAV-DIO-hM3Dq-mCherry/AAV-DIO-mCherry and VTA drug delivery cannula implantation (left). A representative trace showing depolarization of the hM3Dq-mCherry-expressing neuron by CNO (right). **i**, **j**, Effects of chemogenetic activation of VGluT3^DRN^ neurons on the von Frey test in SNI 2W mice (**i**) and SPT in SNI 6W mice (**j**) with drug injection. Significance was assessed by two-way ANOVA followed by Bonferroni’s multiple comparisons test in (**c**, **d**, **f**, **i**, **j**) and two-tailed paired Student’s *t*-test in (**e**). For (**b**), data are shown as box and whisker plots (medians, quartiles (boxes) and ranges minimum to maximum (whiskers); For (**c-j**) data are presented as the mean ± s.e.m. \**P* < 0.05, \*\**P* < 0.01, \*\*\**P* < 0.001, not significant (ns). Details of the statistical analyses are presented in Supplementary Table 1.

To assess the tonic pain, we next tested conditional place preference (CPP), which is correlated with the reward of pain relief. We used this assay to ascertain whether activating VGluT3^DRN→VTA^ terminals in SNI mice induces pain-relief seeking behavior. Mice were tested in a three-chamber CPP assay. Blue light stimulation (10-ms pulse at 20 Hz) was continuously delivered for 30 minutes when mice were restricted in the white chamber, whereas no light were delivered in the black chamber (Fig. 4e). We observed that the ChR2-injected SNI 2W mice displayed preference of the photostimulation-paired chamber. In addition, the ChR2-injected sham control mice also showed CPP, suggesting the involvement of VGluT3^DRN→VTA^ projection in mediating reward (Fig. 4e).

We further examined whether CDB in chronic pain can be modulated by VGluT3^DRN→VTA^ projection which consists co-transmitter of glutamate and 5-HT. We found that selective activation of VGluT3^DRN→VTA^ terminals of SNI 6W mice significantly rescued sucrose preference (Fig. 4f), an indication of alleviated CDB.

## Glutamate is essential for analgesic and anti-depressive effects by the VGluT3^DRN→VTA^ neural excitation

Given co-transmitter of glutamate and 5-HT in the VGluT3^DRN→VTA^ neurons, we hypothesized pain and comorbid depression-modulating VGluT3^DRN^ neurons could act through glutamate and/or 5-HT transmission within the VTA. To test this, AAV-DIO-hM3D(Gq)-mCherry or AAV-DIO-mCherry was injected into the DRN of *VGluT3-Cre* mice and drug delivery cannula was implanted into the VTA (Fig. 4g, h). VGluT3^DRN^ neuronal activation-evoke neuropathic pain relief at SNI 2W and comorbid depression relief at SNI 6W were analyzed following intra-VTA injection of CNQX or ketanserin, selective antagonist of AMPAR and 5-HT2A/2C receptors, respectively. The functional expression of hM3Dq in VGluT3^DRN^ neurons was indicated by neural depolarization by CNO (10 μM) (Fig. 4h). Following i.p CNO (2 mg/kg), the paw withdrawal threshold to von Frey in SNI 2W hM3Dq group was heightened compared to mCherry controls (Fig. 4i), suggesting pain relief by the VGluT3^DRN^ neuronal excitation. Interestingly, the pain alleviation was selectively abolished by intra-VTA injection of CNQX, rather than ketanserin (Fig. 4i). The comorbid depression relief assessed by sucrose preference in SNI 6W mice following VGluT3^DRN^ neuronal excitation was also abrogated by CNQX (Fig. 4j). These data suggest that glutamate plays an essential role in both analgesic and anti-depressive effects by the VGluT3^DRN→VTA^ neurons projection.

## The inactivation of VGluT3^DRN→VTA^ terminals is sufficient to induce hypersensitivity and comorbid depression

The undermined VGluT3^DRN^→DA^VTA^ transmission during the chronic pain prompted us to further examine whether inhibition of VGluT3^DRN→VTA^ terminals would cause pain-like hypersensitivity. We injected *VGluT3-Cre* mice with AAV-DIO-eNpHR-EYFP or AAV-DIO-EYFP into the DRN and implanted optical fibers above the VTA (Fig. 5a, b). Immunostaining and electrophysiological validation confirmed the viability of functional expression of eNpHR in VGluT3^DRN^ neurons (Fig. 5b, c). Mice were then subjected to the von Frey and Hargreaves test during 594-nm yellow light stimulation. We found that optical inhibition of VGluT3^DRN→VTA^ terminals robustly induced both mechanical and thermal hypersensitivity (Fig. 5d, e). Consistently, conditional place avoidance (CPA), an effective assay for measuring ongoing pain, was elicited upon the inactivation of VGluT3^DRN→VTA^ terminals (Fig. 5f). Optogenetic inactivation of VGluT3^DRN→VTA^ projection did not affect the total distance in the open field test (Supplementary Fig. 10c, d), indicating that the motor performance was unchanged.

**Fig 5.**
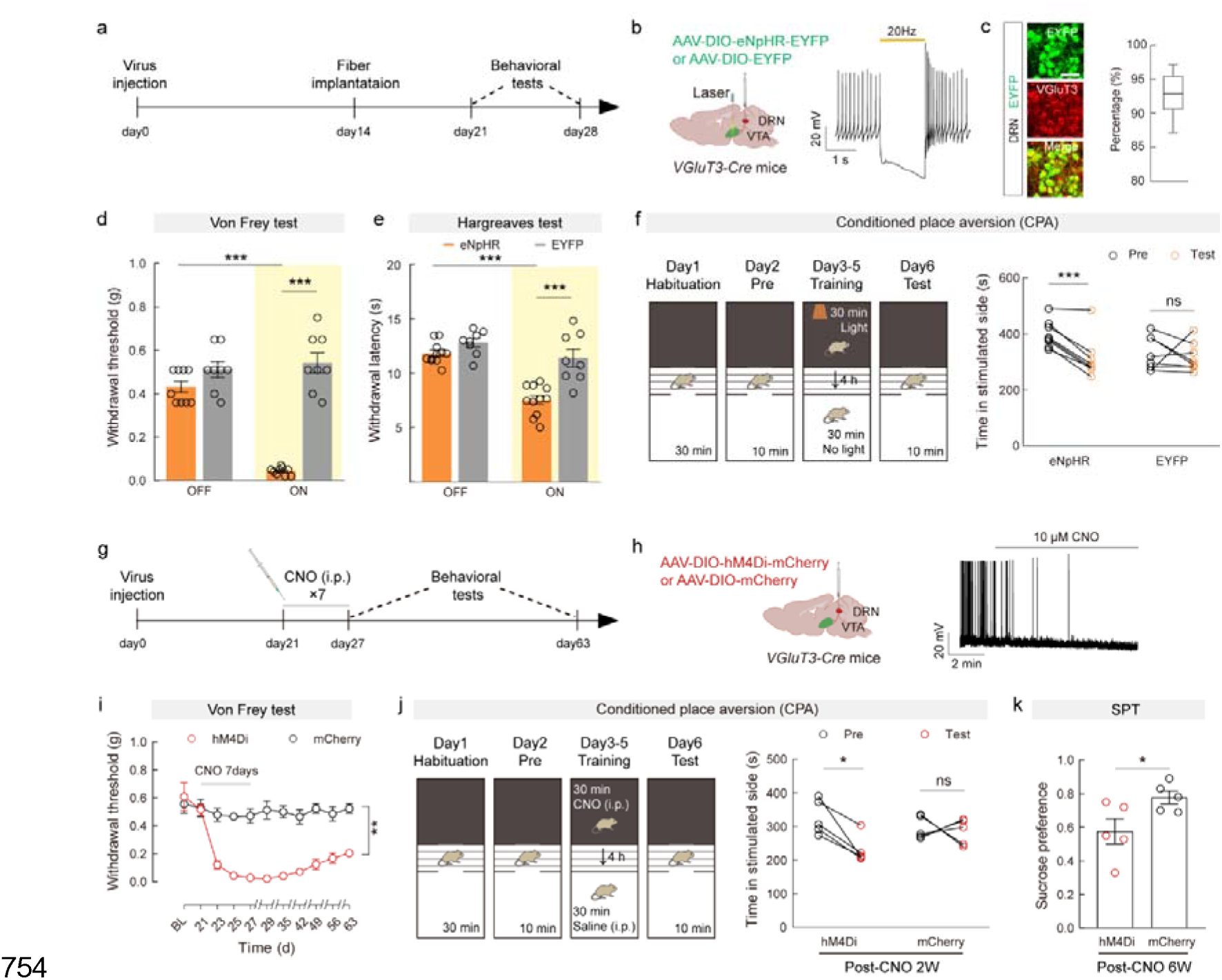
The inactivation of VGluT3^DRN→VTA^ terminals is sufficient to induce hypersensitivity and comorbid depression. **a**, **g**, Schematic of the experimental design. **b**, Schematic of DRN injection of AAV-DIO-eNpHR-EYFP/AAV-DIO-EYFP and VTA optical fiber implantation in *VGluT3*-Cre mice (left). A representative trace of action potential firing of eNpHR-EYFP-expressing neuron during light photostimulation (right). **c**, Representative images (left) and summary data (right) for percentage of EYFP-expressed neurons co-localized with VGluT3 immunofluorescence in the DRN, n = 9 sections from three mice. Scale bar, 50 μm. **d**, **e**, Mechanical paw withdrawal threshold (**d**) and thermal paw withdrawal latency (**e**) of eNpHR-EYFP-expressing and EYFP-expressing mice with (on) or without (off) optogenetic inhibition of VGlut3^DRN→VTA^ terminals. **f**, Experimental design of conditional place aversion (CPA) test (left) and quantification of CPA before (Pre) and after (Test) training of the mice injected with AAV-DIO-eNpHR-EYFP or control AAV-DIO-EYFP (right). **h**, Schematic diagram of DRN injection of AAV-DIO-hM4Di-mCherry/AAV-DIO-mCherry (left) and a representative trace showing hyperpolarization of the hM4Di-mCherry-expressing neuron by CNO (right). **i**, Time-course of mechanical paw withdrawal threshold changes induced by daily injection of CNO for one week in hM4Di-mCherry-expressing and mCherry-expressing mice. **j**, Experimental design of conditional place aversion (CPA) test (left) and quantification of CPA before (Pre) and after (Test) training of the mice injected with AAV-DIO-hM4Di-mCherry or control AAV-DIO-mCherry (right). **k**, Preference for sucrose in the SPT of same animals in **i**, **j**. Significance was assessed by two-way ANOVA followed by Bonferroni’s multiple comparisons test in (**d**, **e**, **i**), two-tailed paired Student’s *t*-test in (**f**, **j**), two-tailed unpaired Student’s *t*-test in (**k**). For (**c**), data are shown as box and whisker plots (medians, quartiles (boxes) and ranges minimum to maximum (whiskers); For (**d-k**) data are presented as the mean ± s.e.m. \**P* < 0.05, \*\**P* < 0.01, \*\*\**P* < 0.001, not significant (ns). Details of the statistical analyses are presented in Supplementary Table 1.

Plastic changes of pain processing circuits caused by sustained nociceptive inputs contribute to pathogenesis of chronic pain^28^. Therefore, we want to know whether prolonged activation of the VGluT3^DRN^ neurons can capture persistent pain as SNI model. To this end, we performed repetitive chemogenetic inhibition of the VGluT3^DRN^ neurons by daily injection of CNO (2 mg/kg) for a week in *VGluT3-Cre* mice with AAV-DIO-hM4Di-mCherry or AAV-DIO-mCherry injection into the DRN (Fig. 5g, h). The hyperpolarization of hM4Di-expressing neurons in the DRN after CNO application confirmed the functionality of the virus (Fig. 5h). As proposed, we observed a persistent mechanical hypersensitivity for more than 6 weeks even after termination of CNO injection in hM4Di group, but not in mCherry group (Fig. 5i). We further tested ongoing pain by CPA paradigm at two days before the first CNO injection. Consistent with above optogenetic inhibition experiments, we found that hM4Di mice showed significant aversion for CNO-paired chamber while mCherry mice spent comparable time before and after CNO treatment (Fig. 5j).

It is noteworthy that hM4Di group displayed a reduction of sucrose preference at six weeks after the first CNO injection (post-CNO 6W) (Fig. 5k), as manifested in SNI 6W mice. These data demonstrate that prolonged silencing of VGluT3^DRN^ neurons mimics both chronic pain-like hypersensitivity and CDB, indicating that tonic activity of VTA-projecting VGluT3^DRN^ neurons is critical for preventing chronic pain-like hypersensitivity and CDB. Collectively, the VGluT3^DRN→^DA^VTA^ circuit modulates chronic neuropathic pain and comorbid depression.

## Medial NAc is the output of the VGluT3^DRN^→DA^VTA^ circuit for neuropathic pain-induced inhibition of DA release

NAc is the major target for the DA^VTA^ neurons processing reward, motivation, and aversive information including pain. We next investigated whether DRN-projected DA^VTA^ neurons innervate the NAc. We injected AAV2/1-Cre into the DRN and Cre-dependent AAV-DIO-mCherry into the VTA of *C57* mice (Supplementary Fig. 7a). Three weeks later, dense projecting terminals were found in both NAcMed and NAcLat, whereas sparse mCherry^+^ fibers were observed in mPFC (Supplementary Fig. 7d). Thus, VTA neurons receiving DRN inputs preferentially project to the NAc but not the mPFC. To examine the functional connectivity of VGluT3^DRN^→DA^VTA^→NAcMed or VGluT3^DRN^→DA^VTA^→NAcLat circuits, we combined optogenetic stimulation of VGluT3^DRN VTA^ inputs with fiber photometry of fluorescence dynamics of DA2m, a G-protein-coupled receptor activation-based DA sensor, in either vNAcMed or NAcLat. To achieve the goal, we injected *VGluT3-Cre* mice with AAV-DIO-ChR2-mCherry into the DRN and AAV-hSyn-DA2m into either NAcMed or NAcLat (Fig. 6a). This allowed us to optostimulate the VGluT3^DRN→VTA^ terminals while simultaneously monitor DA release in the NAcMed or NAcLat using fiber photometry (Fig. 6b). In intact animals, our result revealed instant increment of DA sensor fluorescence both in the NAcMed and NAcLat following 473-nm laser stimulation (2 s, 20 Hz) (Fig. 6c, d), indicating VGluT3^DRN^→DA^VTA^ evoked dopamine release. After nerve injury, intriguingly, we detected an exclusive decrease of dopamine release in the NAcMed upon VGluT3^DRN^ terminals activation (Fig. 6c, d).

**Fig 6.**
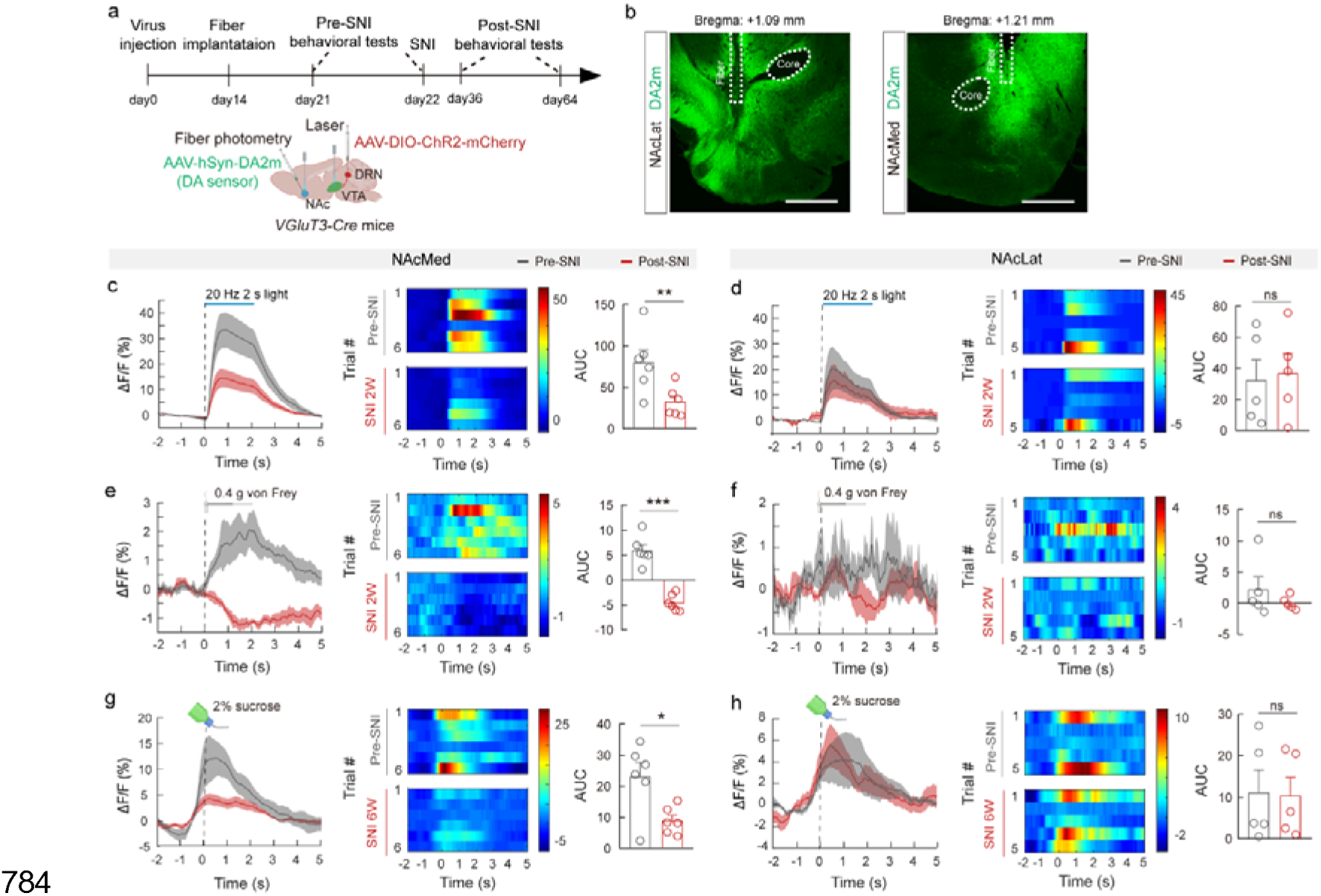
Medial NAc is the output of the VGluT3^DRN^→DA^VTA^ circuit for neuropathic pain-induced inhibition of DA release. **a**, Schematic of the experimental design (top) and schematic diagram of viral injection, VTA laser stimulation and NAc fiber photometry of DA sensor in *VGluT3-Cre* mice (bottom). **b**, Representative images showing optical fiber track for fiber photometry recordings in the NAcMed (left) and NAcLat (right). Scale bars, 500 μm. **c**-**h**, Averaged responses (left), heatmaps (middle) and AUC during 0-5 s (right) showing DA2m signals evoked by optogenetic activation of VGlut3^DRN^ terminals (**c**, **d**), von Frey stimulation (**e**, **f**) in SNI 2W mice compared with pre-SNI mice and sucrose licking in SNI 6W mice compared with pre-SNI mice (**g**, **h**). Significance was assessed by two-tailed paired Student’s *t*-test in (**c**-**h**). All data are presented as the mean ± s.e.m. \**P* < 0.05, \*\**P* < 0.01, \*\*\**P* < 0.001, not significant (ns). Details of the statistical analyses are presented in Supplementary Table 1.

To further explore which sub-region of the NAc is the output of VGluT3^DRN^→DA^VTA^ circuit for neuropathic pain-induced inhibition of DA release, we measured DA release within the NAcMed or NAcLat in responses to 0.4 g von Frey and sucrose consumption before and after SNI. Under physiological condition, von Frey stimulation evoked robust DA release in the NAcMed. In contrast, subtle DA2m fluctuation was observed in the NAcLat (Fig. 6e, f). Sucrose licking-induced DA release was prominent in both NAcMed and NAcLat (Fig. 6g, h). However, von Frey filament and sucrose consumption-induced DA release were both decreased specifically in the NAcMed of SNI 2W and SNI 6W mice, respectively (Fig. 6e, g). Taken together, medial NAc may be the output of the VGluT3^DRN^ DA^VTA^ circuit for neuropathic pain-induced inhibition of DA release.

## Pain and comorbid depression relief by the VGluT3^DRN→VTA^ terminals excitation is via discrete dopaminergic receptors in the NAcMed

D1- and D2-dopamine receptors (D1R and D2R) are two major types of dopaminergic receptors in the NAcMed. We finally investigated the requirement of D1 and/or D2 receptors for neuropathic pain and comorbid depression relief. AAV-DIO-ChR2-mCherry or AAV-DIO-mCherry was delivered into the DRN of *VGluT3-Cre* mice. Two weeks later, mice were subjected to cannula implantation into the NAcMed and SNI surgery (Fig. 7a, b). VGluT3^DRN→VTA^ terminals activation-evoke pain and comorbid depression relief were evaluated by intra-NAcMed injection of SCH23390 or Eticlopride, selective antagonist of D1R and D2R, respectively. Following 473-nm laser stimulation of VGluT3^DRN→VTA^ terminals, the paw withdrawal threshold to von Frey was elevated in ACSF-perfused SNI 2W mice (Fig. 7c). However, the elevation was selectively eliminated by Eticlopride, but not SCH23390 (Fig. 7c). In contrast, VGluT3^DRN→VTA^ terminals activation-evoked comorbid depression relief in SNI 6W mice was absent in SCH23390-treated group, but not in Eticlopride-treated mice (Fig. 7d). Of note, the neuropathic pain and comorbid depression relief was not affected by NAcLat injection of either antagonist (Fig. 7e, f). Overall, our study reveals that a glutamatergic circuit from VGluT3^DRN^ to DA^VTA^ undergoes maladaptive changes during neuropathic pain, and VGluT3^DRN^→DA^VTA^ circuit modulates pain-like hypersensitivity and CDB via D2R and D1R in NAcMed, respectively (Fig. 7g).

**Fig 7.**
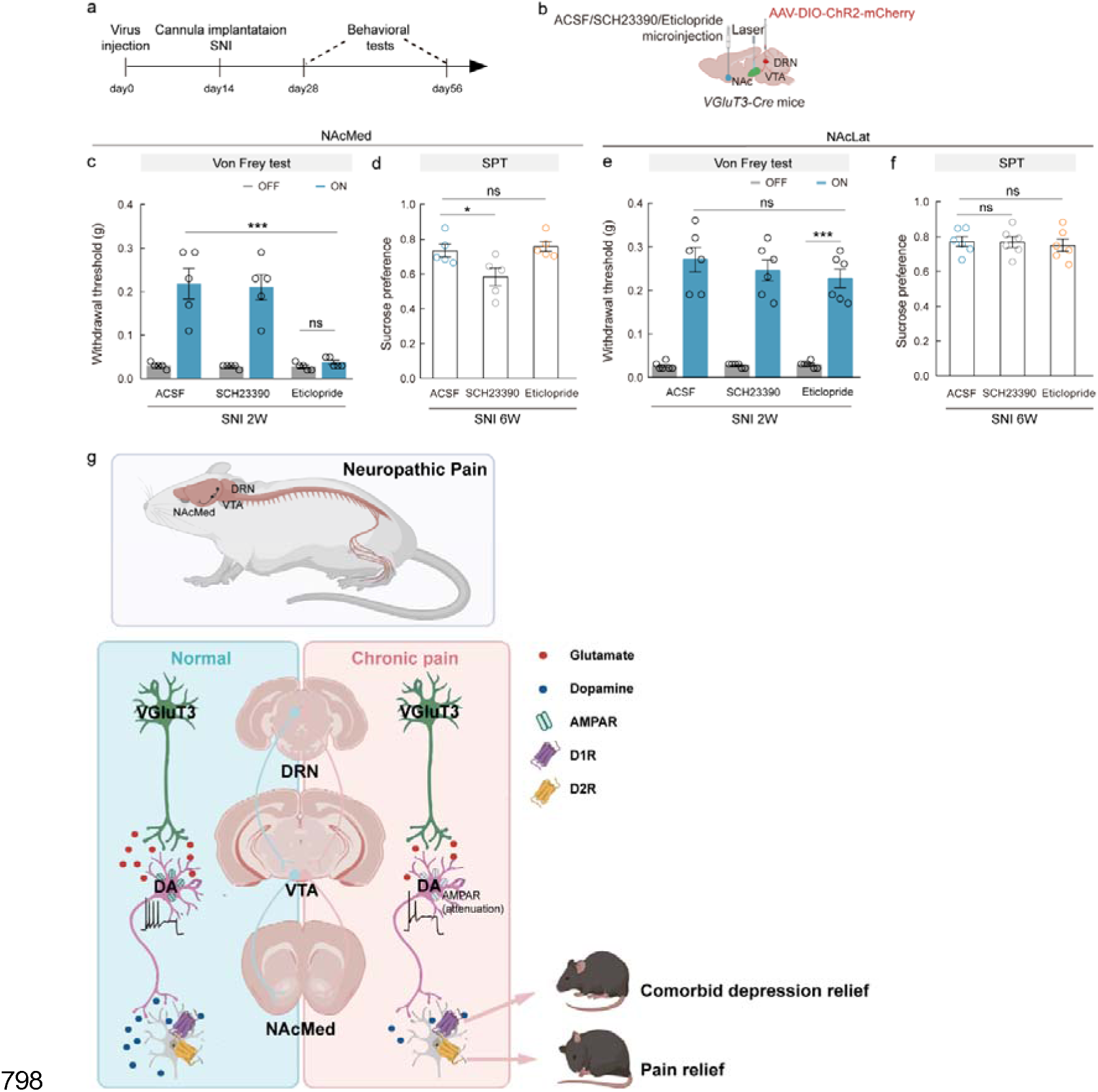
D1- and D2-dopamine receptors contribute to analgesia and anti-depressive effects of VGluT3^DRN^→DA^VTA^ circuit. **a**, Schematic of the experimental design. **b**, Schematic diagram of DRN viral injection, VTA laser stimulation and NAc drug delivery cannula implantation. **c**-**f**, Effects of optogenetic activation of VGluT3^DRN^ terminals on the von Frey test and SPT with drug infusion into the NAcMed (**c**, **d**) and NAcLat (**e**, **f**). **g**, Schematic summary of maladaptive changes of VGluT3^DRN^→DA^VTA^→D1/D2^NacMed^ circuit during neuropathic pain. Significance was assessed by two-way ANOVA followed by Bonferroni’s multiple comparisons test in (**c**, **e**), and one-way ANOVA followed by Bonferroni’s multiple comparisons test in (**d**, **f**). All data are presented as the mean ± s.e.m. \**P* < 0.05, \*\*\**P* < 0.001, not significant (ns). Details of the statistical analyses are presented in Supplementary Table 1.

## Discussion

Mutual deterioration of chronic pain and comorbid depressive symptom brings great challenge for effective clinical treatment. In the present study (Fig. 7g), we identified a crucial role of VGluT3^DRN^→DA^VTA^→D2/D1^NAcMed^ circuit in establishing and modulating chronic neuropathic pain and comorbid depression. We reported that chronic pain dampens VGluT3^DRN^→DA^VTA^ glutamatergic transmission and consequently decreases DA release in specific subregion of the NAc shell. In addition, VGluT3^DRN^→DA^VTA^ circuit activation relieves pain-like hypersensitivity and CDB via discrete dopamine receptors in the NAcMed. VGluT3^DRN^→DA^VTA^ circuit inactivation is sufficient to drive chronic pain-like hypersensitivity and depression-like behavior. These findings provide a novel neural substrate for treating chronic pain and comorbid depression.

It has long been recognized that a large proportion of VTA DA neurons is inhibited by acute noxious stimuli^9,17,18^, and circuit-based GABAergic inhibition has been involved in acute pain-induced DA^VTA^ neural hypoexcitability^9^. However, the maladaptive and dynamic changes of DA neurons under chronic pain and comorbid depression remained elusive. Moreover, circuit mechanisms underpinning chronic pain-induced DA neural dysfunction are unclear. In the current work, we demonstrated that chronic pain decreases excitability of DA^VTA^ neurons, and dampened activity of the VGluT3^DRN^→DA^VTA^ pathway might serve as one of the mechanisms (Fig. 1 and 3).

Our results discovered a dynamic stage-dependent down-regulation of *Ih*, which is mediated by HCN channels that predominantly distribute at the dendrites of DA^VTA^ neurons^29^ (Fig. 1j and 3q), and plays a pivotal role in temporal integration of synaptic inputs^23^. The undermined *Ih* would lower the synchrony of neuronal networks from a wide range of synaptic inputs and adversely affect postsynaptic neuronal function^23^. Otherwise, lines of preclinical studies showed that enhancing *Ih* exerts antidepressant effect or achieves homeostatic resilience^30–32^. Thus, our findings reasoned a causal connection between the pathophysiological decreases in *Ih* with stage-dependent CDB manifestation during neuropathic pain.

The neural activity of DA^VTA^ neurons are set by both intrinsic electrophysiological properties and sophisticated excitatory/inhibitory synaptic input balance^33^. Our results provided evidence that reduced excitatory drive from presynaptic VGluT3^DRN^ neurons plays a role. The DRN is composed of molecularly and functionally heterogeneous neural subpopulations^34,35^. The role of DRN serotonergic system in descending pain modulation and relieving depression and several major psychiatric disorders has been well studied^36^. However, how adaptation of the DRN→VTA circuit, which is involved in signaling reward^13,26^, may contribute to neuropathic pain and CDB is unknown. Viral-genetic tracing revealed that VTA-projecting DRN neurons consisted of a large population (∼59%) of VGluT3^+^ neurons. Notably, ∼23% within them were dual VGluT3-serotonin neurons (Fig. 2g). The functional glutamatergic and serotonergic synapse between VGluT3^DRN^ neurons and DA^VTA^ neurons was evidenced by electrophysiology. These findings are consistent with previous studies showing that a subset of DRN neurons projecting to VTA co-release glutamate and serotonin^13,26,37^. Pathophysiologically, SNI decreased the population activity of VGluT3^DRN→VTA^ afferents in responses to pain stimuli and sucrose licking, and weakened VGluT3^DRN^→DA^VTA^ synaptic connectivity (Fig. 2). Moreover, reduced excitability and a stage-dependent *Ih* down-regulation of DRN-targeted DA^VTA^ neurons were detected (Fig. 2). Behaviorally, bidirectional manipulation of VGluT3^DRN^→DA^VTA^ circuit could alleviate or mimic the SNI-induced pain and depression-like behavior (Fig. 4 and 5). These evidences collectively establish a compelling correlation between aberrant activity of VGluT3^DRN^→DA^VTA^ circuit and SNI-induced pain and CDB. Of note, glutamate from VGluT3^DRN→VTA^ terminals underlies the analgesia and antidepressant effects. It remains to be addressed the source of glutamate from which subpopulation of VGluT3-expressing neurons containing VGluT3-only or dual VGluT3-serotonin.

Although 5-HT in VGluT3^DRN^→VTA circuit exerts no influence on CDB in our study, the serotonergic adaptation of DRN→CeA pathway in neuropathic pain has been implicated in comorbid depression^15^. In other words, DRN neurons could regulate comorbid depression in neuropathic pain via different transmitters acting through distinct downstream brain areas. This insight offers hope for circuit-based specific receptors manipulation towards chronic pain and CDB treatment. In addition, the retrograde trans-synaptic projections of DRN neurons included the spinal cord, locus coeruleus, periaqueductal gray and basolateral amygdaloid nucleus (Supplementary Fig. 11), which had been involved in processing pain information^38,39^. How these DRN-centered circuits cooperate with each other and how these circuits undergo adaptations are worth further study, providing a more detailed understanding of the pathogenesis of chronic pain and CDB.

Dopamine neurotransmission in the nucleus accumben is commonly considered a primary regulator of motivational drive and a critical target of drug abuse^16,40,41^. Human studies also demonstrated promising efficacy of NAc DBS in treatment-refractory pain and major depression^42,43^, yet the precise NAc subregions for analgesia and antidepressant effects are ambiguous, especially for comorbid depression in neuropathic pain. In our study, while we observed that DRN-downstream VTA neurons send dense innervation to both NAcLat and NAcMed, neuropathic pain evoked region-specific DA alteration was preferentially found in the NAcMed. Moreover, D2 and D1 receptors in the NAcMed underlay the pain and CDB relief by VGluT3^DRN→VTA^ terminals activation. These findings advance the understanding of how chronic pain alters subregion-specific dopaminergic system and behaviors (Fig. 6 and 7). Although D1 and D2 receptors in NAc have long been implicated in reward and aversive behaviors, respectively^44^, a recent study reported that activation of D2-dopamine receptor-expressing striatal medium spiny neurons (D2-MSNs) in NAc could also drive reinforcement^45^. Therefore, the role of D1-dopamine receptor-expressing MSNs (D1-MSNs) and D2-MSNs in mediating positive and negative motivational valence need to be revisited. The functional divergence between D1-MSNs and D2-MSNs could arise from their different neural circuit connectivity^46,47^. Future studies on input-output circuits of the two neuronal populations are needed to help revealing their roles in different chronic pain-related behaviors.

Overall, our study demonstrated the dysfunctional adaptation of the glutamatergic VGluT3^DRN^→DA^VTA^ pathway during chronic pain and CDB development. Importantly, we revealed subregion-specific alteration in DA release and discrete dopaminergic receptors for managing chronic pain and CDB, and provided important mechanistic insight for the transition from sensory to affective component of chronic pain, which may shed light on identifying potential therapeutic targets for treating chronic pain and CDB in humans.

## Acknowledgments

We thank Dr. Qiufu Ma in Westlake University for critical comments on the manuscript. This study was supported by the National Natural Science Foundation of China (grants 32070999, 32271048 and U20A20357), Anhui Provincial Natural Science Foundation (grant 2008085J16), and the Fundamental Research Funds for the Central Universities (WK2070210004 and WK9110000056).

## Author contributions

X. Y. W. and W. B. J. designed the studies and conducted most of the experiments and data analysis. X. Y. W. wrote the first draft. X. X., R. C., L. B. W. and X. J. S generated some molecular and behavioral data. P. F. X. and X. Q. L. managed the mouse colonies used in this study. J. W. and Y. Y. L. were involved in the overall design of the study. Z. Z., X. F. L. and Y. Z. were involved in the overall design of the project, data analysis, and editing of the final manuscript.

## Declaration of interests

The authors declare no competing interests.

All data necessary to understand and assess the conclusions of this study are available in the main text or the supplementary materials. There are no restrictions on data availability in the manuscript.

## Methods

### Animals

In all experiments, male and female mice aged 8-10 weeks were used. *C57BL/6J* mice were purchased from Beijing Vital River Laboratory Animal Technology, *VGluT3-Cre* and *Ai14 (RCL-tdTOM)* mice were purchased from Jackson Laboratories, *DAT-Cre* and *GAD2-Cre* mice were gifts from GQ Bi and Z Zhang, respectively. Mice were kept on a 12-h light/dark cycle (lights on at 7am) at a stable temperature of 23 ± 1 °C. Food and water were freely available. All animal protocols were approved by the Animal Care and Use Committee of the University of Science and Technology of China.

### Chronic pain model

Chronic neuropathic pain was induced in mice by SNI surgery. Under anesthesia with isoflurane, skin incision was made on the left thigh and the muscle was gently separated to explore the sciatic nerve consisting of the sural, common peroneal and tibial nerves. The common peroneal and tibial nerves were separated from the sural nerves, tightly ligated with a nonabsorbent 4-0 chromic gut sutures (Ethicon) and transected together. About 2 mm sections from the nerves were removed. The skin was stitched and disinfected with iodophor. For the sham group, mice were managed in the same manner, but the nerves were not ligated.

### Stereotactic surgeries

Under anesthesia induced with an intraperitoneal injection of pentobarbital (20 mg/kg), the mice were fixed in a stereotactic frame (RWD) with a heating pad. Ophthalmic ointment was applied to avoid corneal drying. An incision was made along the midline to expose the skull surface. A volume of 100-300 nl virus was injected in the target region at a rate of 30 nl/min using a pulled glass micropipette connected to a microsyringe pump (KD Scientific). The pipette was stayed for additional 5 min after the injection and then slowly withdrawn.

For optogenetic manipulation of DRN-VTA circuit, we injected rAAV2/9-Ef1α-DIO-hChR2(H134R)-mCherry-WPRE-pA (AAV-DIO-ChR2-mCherry, 4.50E+12 vg/ml, 200 nl), rAAV2/9-Ef1α-DIO-eNpHR3.0-EYFP-WPRE-pA (AAV-DIO-eNpHR-EYFP, 5.36E+12 vg/ml, 200nl) into DRN (anterior-posterior (AP): -4.50 mm; medial-lateral (ML): 0 mm; dorsal-ventral (DV): -3.15 mm) of *VGluT3-Cre* mice. For chemogenetic behavioral experiments, the rAAV2/9-Ef1α-DIO-hM3D(Gq)-mCherry-WPRE-pA (AAV-DIO-hM3Dq-mCherry, 5.27E+12 vg/ml, 200nl), the rAAV2/9-Ef1α-DIO-hM4D(Gi)-mCherry-WPRE-pA (AAV-DIO-hM4Di-mCherry, 5.18E+12 vg/ml, 200 nl) viruses were injected into the DRN of *VGluT3-Cre* mice and the VTA (AP, -1.18 mm; ML, -2.65 mm; DV, -4.25 mm) of *DAT-Cre* mice. The rAAV2/9-Ef1α-DIO-mCherry-WPRE-pA (AAV-DIO-mCherry, 5.14E+12 vg/ml, 200 nl), rAAV2/9-DIO-EYFP-WPRE-pA (AAV-DIO-EYFP, 5.14E+12 vg/ml, 200 nl) were used as the controls.

For retrograde monosynaptic tracing, helper viruses that contained rAAV2/9-Ef1α-DIO-RVG-WPRE-pA (AAV-DIO-RVG, 5.29E+12 vg/ml) and rAAV2/9-Ef1α-DIO-EGFP-2a-TVA-WPRE-pA (AAV-DIO-TVA-GFP, 5.56E+12 vg/ml, 1:2, 200 nl) were co-injected into the right VTA of *DAT-Cre* mice or *GAD2-Cre* mice. Three weeks later, the rabies virus RV-ENVA-ΔG-DsRed (2.0E+08TU/ml, 200 nl) was injected into the same site in the VTA. Seven days after the last injection, mice were perfused, and brain slices were prepared for retrograde monosynaptic tracing. For fluorogold retrograde tracing, *VGluT3-tdTOM* mice were injected with retrograde tracer Fluorogold (FG; diluted with saline, 1%, 300 nl) into the VTA. Seven days after injection, brain slices were co-staining with TPH2-specific antibodies to distinguish phenotypes among DRN neurons projecting to the VTA. For retrograde trans-synaptic tracing experiments, PRV-CAG-EGFP (2.00E+09 PFU/ml, 150 nl) was injected into the DRN of *C57* mice. Four days after PRV injection, mice were killed to trace EGFP signal.

For monosynaptic anterograde tracing, rAAV2/1-hSyn-CRE-WPRE-hGH-pA (AAV2/1-Cre, 1.00E+12 vg/mL, 200 nl) was injected into the DRN of *C57* mice to allow the virus to express Cre in the downstream soma. Simultaneously, AAV-DIO-mCherry (200 nl) was injected into the VTA. After 3 weeks expression, brain slices were prepared for triple tracing or co-staining with TH-specific antibodies or GABA-specific antibodies in the VTA. Unless otherwise stated, all viruses were packaged by BrainVTA.

### *In vivo* optogenetic manipulations

Two weeks after virus injection, an optical fiber (200 μm, NA = 0.37; Inper) was unilaterally implanted into the right VTA (AP: -3.20 mm, ML: -0.45 mm, DV: -3.80 mm) of the mice fixed in a stereotactic frame with a heating pad. The delivery of blue light (473 nm, 3-5 mW, 10 ms duration at 20 Hz) or yellow light (594 nm, 8-10 mW, 10 ms duration at 20Hz) was controlled using a ThinkerTech stimulator. An identical stimulus protocol was applied in the control group. By examining the location of the fibers, only mice with the correct locations of optical fibers and viral expression were used for data analysis.

### Fiber photometry

To record calcium fluorescence of VTA DAergic neurons, rAAV2/9-Ef1α-DIO-GCaMP6m-WPRE-pA (AAV-DIO-GCaMP6m, 6.21E+12 vg/ml, 300 nl) was unilaterally injected into the right VTA (AP: -3.20 mm, ML: -0.45 mm, DV: -4.25 mm) of *DAT-Cre* mice. Two weeks after virus injection, an optic fiber (200 μm, 0.37 NA; Inper) was placed into the right VTA (AP: -3.20 mm, ML: -0.45 mm, DV: -3.95 mm). To record calcium fluorescence of VGluT3^DRN^ terminals in the VTA, 300 nl of AAV-DIO-GCaMP6m was injected into the DRN of *VGluT3-Cre* mice. After allowing for 14 days of virus expression, an optical fiber (200 μm, 0.37 NA; Inper) was implanted into the right VTA. To measure dopamine release, rAAV2/9-hSyn-DA4.4 (AAV-hSyn-DA2m, 5.60E+12 vg/ml, 300 nl) was injected into the NAcMed (AP: +1.30 mm, ML: -0.75 mm, DV: -4.50 mm) and NAcLat (AP: +1.0 mm, ML: -1.80 mm, DV: -4.90 mm) of ChR2-expressed *VGluT3-Cre* mice, and optical fibers were implanted over the NAcMed (AP: +1.30 mm, ML: -0.75 mm, DV: -4.20 mm) and NAcLat (AP: +1.0 mm, ML: -1.80 mm, DV: -4.60 mm) to record change of fluorescence. Photometric recordings were conducted using the fiber photometry recording system (ThinkerTech) 7 days after the fibers-implantation procedures to ensure adequate animal recovery.

Calcium-dependent fluorescence signals were obtained by stimulating neurons and terminals expressing GCaMP6m with laser intensities for the 470 nm wavelength bands (40 μW), and 410 nm signal (20 μW) was further used to correct for movement artifacts. Light emission was recorded using a sCMOS Camera, and the values of Ca^2+^ signal changes (ΔF/F) by calculating (F−F0)/F0 (Averaged baseline fluorescence signal recorded) were analyzed by MATLAB. A 5-s window around stimulation point was analyzed, with the period 2 s before stimulus onset taken as baseline. Ca^2+^ responses during the first six times of required behavior of each mouse were analyzed. For fiber photometry recordings in von Frey test, the filament stimulation (0.4 g) was delivered onto the ipsilateral hindpaws (nerve-injured side) after 30 min habituation, Ca^2+^ responses evoked by von Frey stimulation in SNI 2W mice compared with pre-SNI mice were recorded. For sucrose licking test, mice were habituated with a bottle of 1% sucrose for 24 h followed with water deprivation for 24 h, then were given free access to the bottle of 1% sucrose, Ca^2+^ responses evoked by sucrose licking in SNI 2W/6W mice compared with pre-SNI mice were recorded. For optogenetic activation test, optical stimulation (473 nm, 3-5 mW, 10 ms duration at 20 Hz) was delivered for 2 s through the fiber implanted in VTA. The fluorescence signals were recorded to measure dopamine release in SNI 2W mice compared with pre-SNI mice.

### *In vivo* pharmacological approach

After allowing for 14 days of virus expression, a guide cannula (0.34 mm, RWD) was implanted into areas of interest, which included the VTA (AP: -3.20 mm, ML: -0.45 mm, DV: -3.85 mm), NAcMed (AP: +1.30 mm, ML: -0.75 mm, DV: -4.10 mm) and NAcLat (AP: +1.0 mm, ML: -1.80 mm, DV: -4.50 mm) of the mice fixed in a stereotactic frame. The implant was secured to the skull of the animal with dental cement, and mice were allowed to recover from surgery over 7 days before subsequent behavioral experiments. Microinjections were administered 30 min before testing, and antagonist dissolved in ACSF (200 nl) was microinjected at a rate of 200 nl/min. To test the participation of glutamatergic or serotonic receptors within VTA in pain and depression relief, we administered microinjections of ACSF, glutamate AMPA receptor antagonist CNQX (1 μg) or serotonin 5-HT2a and 5-HT2c receptor antagonist ketanserin (1 μg) in the VTA. To evaluate the participation of D1 or D2 receptors within NAc in pain and depression relief, we administered microinjections of ACSF, D1 receptor antagonist SCH23390 (0.1 μg) or D2 receptor antagonist eticlopride (1 μg) in the NAcMed and NAcLat.

### Immunofluorescence and imaging

Mice anesthetized with isoflurane were transcardially perfused with 0.1 M PBS and 4% paraformaldehyde in PBS. Brains were removed and post-fixed in 4% paraformaldehyde at 4 °C overnight, dehydrated in 20% and then 30% sucrose until they sank. Coronal sections (35 μm) were sliced on a cryostat (Leica CM1950). For immunofluorescence, sections were washed with PBS three times (5 min each) and were incubated with blocking buffer (0.3% Triton X-100, 10% goat serum in PBS) for 1 h at room temperature, then they were incubated (12-24 h at 4 °C) with the primary antibodies anti-TPH2 (1:500, rabbit, Abcam), anti-Vglut3 (1:500, rabbit, Synaptic Systems), anti-GABA (1:500, rabbit, Sigma), anti-TH (1:1000, rabbit, Emd millipore). After rinsing with PBS, the sections were incubated with the corresponding fluorophore-conjugated secondary antibodies (1:500, Jackson) for 2 h at room temperature. Sections were then washed three times with PBS, stained with DAPI, and coverslipped with fluorescent mounting medium. Confocal images were captured on an Olympus FV3000 microscope and analyzed with ImageJ software.

### *In vitro* electrophysiological recordings

For slices preparation, the mice were deeply anesthetized with isoflurane, and intracardially perfused with ice-cold oxygenated N-Methyl-D-glucamine (NMDG) artificial cerebrospinal fluid (ACSF) that contained (in mM) 93 NMDG, 2.5 KCl, 30 NaHCO3, 1.2 NaH2PO4, 20 HEPES, 25 glucose, 2 thiourea, 3 Na-pyruvate, 0.5 CaCl2, 5 Na-ascorbate, 10 MgSO4 and 3 glutathione (PH 7.3-7.4, 300-305 mOsm). Coronal slices (300 μm) were then removed to ice-cold oxygenated NMDG-ACSF and sectioned on a vibrating microtome (VT1200s, Leica). The brain slices were allowed to recover in oxygenated NMDG-ACSF for 10 min at 32 °C, followed by oxygenated N-2-hydroxyethylpiperazine-N-2-ethanesulfonic acid (HEPES) ACSF that contained (in mM) 92 NaCl, 2.5 KCl, 1.2 NaH2PO4, 3 Na-pyruvate, 30 NaHCO3, 20 HEPES, 25 glucose, 2 thiourea, 5 Na-ascorbate, 2 CaCl2, 2 MgSO4 and 3 GSH (PH 7.3-7.4, 300-305 mOsm) for more than 1 h at 25 °C. Slices were transferred to the recording chamber for electrophysiological recordings. Recordings were performed in oxygenated standard ACSF that contained (in mM) 3 HEPES, 129 NaCl, 1.2 KH2PO4, 3 KCl, 2.4 CaCl2, 1.3 MgSO4, 10 glucose, and 20 NaHCO3 (PH 7.3-7.4, 300-310 mOsm, oxygenated with 95% O2 and 5% CO2) at 32 °C.

Patch-clamp electrophysiology data were analyzed with Clampfit pClamp 10.0 software using MultiClamp 700B (Molecular Devices) and Digidata 1550B (Molecular Devices), digitized at 5 kHz, and filtered at 2 kHz. The pipette (6-8 MΩ) was pulled by a micropipette puller (P-1000, Sutter instrument) and filled with the internal solution that contained (in mM): 130 K-gluconate, 5 KCl, 4 Na2ATP, 0.5 NaGTP, 20 HEPES, 0.5 EGTA, (PH 7.28, 290-300 mOsm). To measure *I*h currents, neurons were voltage-clamped at -70 mV and stepped to -120 mV in increments of 10 mV. To record an input–output curve of neuronal excitability, 500 ms pulses with 10 pA command current steps were injected from -60 to +150 pA, and the numbers of spikes were quantified for each step. The rheobase was defined as minimal current required to evoke an action potential.

To examine the monosynaptic nature of eEPSCs and eIPSCs in VTA neurons following light activation of VGluT3^DRN^ inputs, neurons were held at -70 mV and 0 mV, respectively. TTX (1 μM) was used to block action potential-based synaptic transmission. Both TTX (1 μM) and 4-AP (100 μM) were used to restore monosynaptic current. The response jitter was calculated by measuring the standard deviation of the latency values of consecutive EPSCs for VTA neurons. For valuating synaptic identities, AMPA-mediated fast EPSCs were blocked by the bath application of CNQX (10 μM), and repetitive light stimulation (20 s; 20 H)-evoked slow IPSP was largely abolished by the addition of the 5-HT receptor antagonist ketanserin (10 μM). For evaluating presynaptic mechanism, paired pulses (10 ms duration) with interval of 50 ms (ISI 50 ms) were delivered, and the PPR was calculated as the amplitude ratio EPSC2/EPSC1.

To measure the AMPA/NMDA current ratio, the pipettes were filled with intracellular solution that contained (in mM) 135 Cs-Meth, 10 KCl, 1 MgCl2, 2 QX-314, 0.3 GTP-Na, 4 ATP-Mg, 0.2 EGTA, and 20 phosphocreatine, (PH 7.28, 290-300 mOsm). Cells were first clamped at -70 mV, and the AMPA receptor-mediated EPSCs were recorded. To record the NMDA receptor-mediated EPSCs, cells were clamped at +40 mV in the presence of bicuculline, and EPSCs were determined 50 ms after the peak of the current response.

### SPT

Mice were habituated with two identical bottles of 1% sucrose for 48 h followed with 48 h of water individually, then subjected to water deprivation for 24 h. For the test day, the mice were given access to a two-bottle choice for 2 h, one containing water and the other containing 1% sucrose, and the bottles positions were switched after 1 h. Sucrose preference was calculated as the proportion of 1% sucrose in the total liquid consumed. For the optogenetic experiment, optostimulation (30 min on/30 min off/30 min on/30 min off) was applied during test day. For chemogenetic experiments, a single injection of CNO (2 mg per kg) was given 30 min before the behavior tests.

### OFT

The open field arena was a square box (40 × 40 cm) within a sound-attenuated room. Mice were allowed to freely explore their surroundings, and the total distance traveled was measured by Smart3 software. To assess the effect of optogenetic activation or inhibition on locomotor activity, mice were tested for a 5-min session.

### Von Frey test

The von Frey filaments test was used to assess the onset and maintenance of mechanical allodynia. Mice were individually placed in a transparent plastic box (6.5 cm × 6.5 cm × 6 cm) to habituate about 30 min for 3 continuous days before the behavior tests. The von Frey filaments (bending force ranging from 0.02 to 2 g, North Coast Medical), which increased in stimulation value, quantified mechanical allodynia by measuring the hind paw response. A positive response included brisk paw withdrawal, flinching, licking or shaking. Using Dixon’s up-down metho^48^, the stimulus producing 50% likelihood of a positive response was determined and taken as the paw withdrawal threshold. For mechanical withdrawal frequency test in Extended Data Fig 2a, filaments were applied to the lateral region of the hindpaws. Withdrawal frequencies were recorded (5 applications per filament, each applied 30 s apart).

### Hargreaves test

Thermal withdrawal latency was measured with Hargreave’s Apparatus (Model390; IITC Life Science Inc, Woodland Hills, CA). Mice were placed individually in chambers on top of a glass platform, and allowed to habituate for at least 60 min before testing. A radiant heat beam was focused onto the ipsilateral hindpaws (nerve-injured side), and paw withdrawal latency was recorded with 5 trials per animal. Five trials were conducted with a minimal interval of 5 minutes. The maximum and minimum PWL were excluded to minimize the variation, and average of the remaining 3 trials was calculated for each mouse. A cut-off latency of 20 s was set to avoid tissue damage.

### RTPA test

2 weeks after SNI surgery, the mice were placed in a custom-made chamber with two distinct visual compartments. The mice moved freely between two chambers for 10 min (Pre). Then we assigned one side as the filament (0.4 g) stimulation side for the following 10-min test. Initially the mouse was placed in the non-stimulated side of the chamber, and every time the mouse crossed to the stimulation side, the filament stimulation was delivered until the mouse crossed back into the non-stimulation side. In the post-stimulation phase, the mouse freely explored two chambers again. The locations of mice were tracked by a video camera positioned above the chamber.

### CPP test

The conditioned place-preference test consisted of 6 d and was performed in the box consisting of two unique conditioning chambers with a neutral middle chamber. On day 1 (Habituation), the mice were habituated to the box for 30 min. On days 2 (Pre), individual mice were placed in the neutral chamber and allowed to freely explore the entire box for 10 min, then showed a small preference for one of the two chambers. On days 3-5 (Training), mice were confined to preferred chambers for 30 min without light. Four hours after the conditioning (No light), mice were individually conditioned in the non-preferred chamber for 30 minutes with blue light stimulation. On day 6 (Test), like day 2, mice were placed in the middle chamber and allowed to explore the entire box for 10 min. The time spent in non-preferred chamber was analyzed to determine whether optogenetic activation was able to produce CPP.

### CPA test

To test the roles of optogenetic inhibition of VGluT3^DRN→VTA^ terminals in aversive aspects, the same procedures were followed as the CPP assay, except that the preferred chamber was assigned as yellow light-paired side and the time spent in preferred chamber was analyzed. Similarly, during CPA assay for chemogenetic inhibition of VGluT3^DRN^ neurons, the preferred chamber was re-designated as CNO-paired chamber, in which the animal was injected with CNO (2 mg per kg).

### Quantification and statistical analysis

Data were analyzed with GraphPad Prism v.8.0.1, Olympus FV10-ASW 4.0a Viewer, ImageJ, Clampfit software v.10.0, and MATLAB software. All experiments and data analyses were conducted double-blind, including fiber photometry, immunofluorescence, electrophysiology and behavioral analyses. No data were excluded from the analyses. Experimental and control animals were randomized throughout the study. Student’s *t*-tests (paired and unpaired) and one-way or two-way ANOVA tests followed by Bonferroni’s test for multiple comparisons were used to determine statistical differences. All data are presented as the mean ± s.e.m. Statistical results in the figures are presented as symbols \**P* < 0.05, \*\**P* < 0.01, \*\*\**P* < 0.001.

---

**Supplementary Figure 1.**
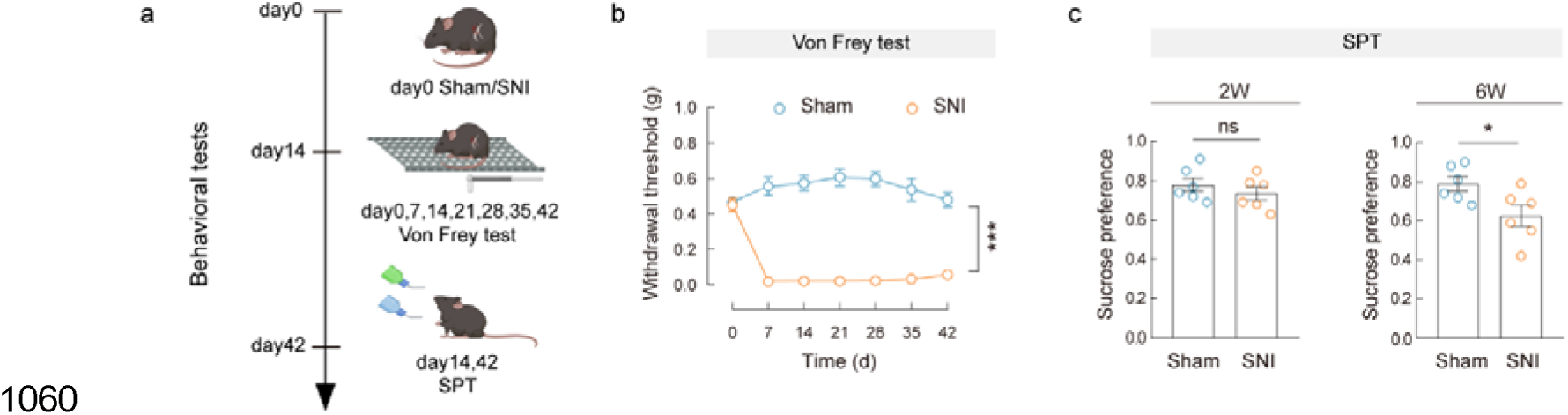
Time course of SNI-induced mechanical hypersensitivity and comorbid depressive-like behavior. **a**, Schematic of the experimental design. **b**, **c**, Performance of mice treated with sham or SNI in von Frey test (b) and SPT (c). Significance was assessed by two-way ANOVA followed by Bonferroni’s multiple comparisons test in (**b**), and two-tailed unpaired Student’s *t*-test in (**c**). All data are presented as the mean ± s.e.m. \**P* < 0.05, \*\*\**P* < 0.001, not significant (ns). Details of the statistical analyses are presented in Supplementary Table 1.

**Supplementary Figure 2.**
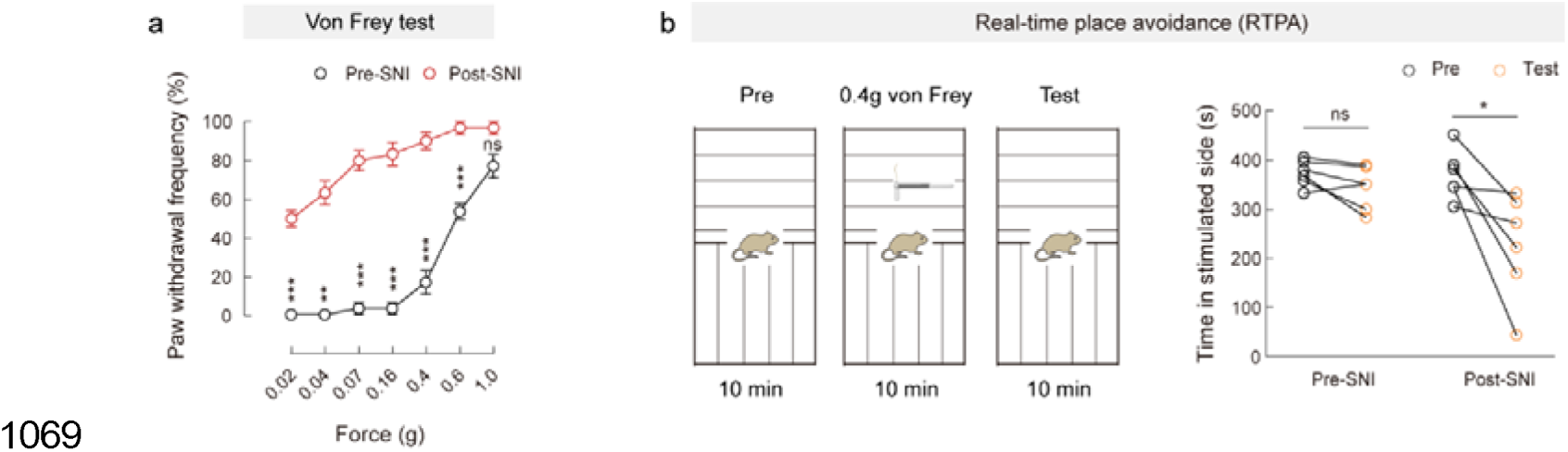
Mechanical withdrawal frequency to von Frey filament and RTPA test before and after SNI. **a**, Statistical data for mechanical withdrawal frequencies to von Frey filaments applied on the hind paw in *C57* mice before (Pre-SNI) and after (Post-SNI 2W) SNI. **b**, Experimental design of real-time place avoidance (RTPA) test (left) and quantification of RTPA before (Pre) and after (Test) training of the Pre-SNI or Post-SNI 2W mice (right). Significance was assessed by two-way ANOVA followed by Bonferroni’s multiple comparisons test in (**a**) and two-tailed paired Student’s *t*-test in (**b**). All data are presented as the mean ± s.e.m. \**P* < 0.05, \*\**P* < 0.01, \*\*\**P* < 0.001, not significant (ns). Details of the statistical analyses are presented in Supplementary Table 1.

**Supplementary Figure 3.**
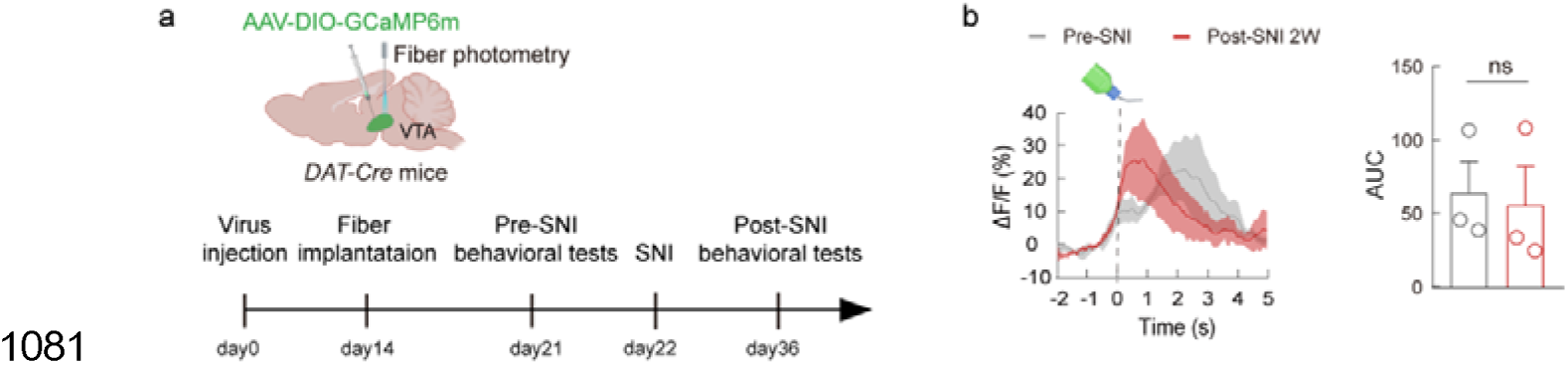
Ca2^+^ signal of DA^VTA^ neurons in *DAT-Cre* mice. **a**, Schematic of the experimental design and schematic diagram of fiber photometry of DA^VTA^neurons. **b**, Averaged responses (left) and AUC during 0-5 s (right) showing Ca^2+^ responses evoked by sucrose licking in SNI 2W mice compared with pre-SNI mice. Significance was assessed by two-tailed paired Student’s *t*-test in (**b**). Not significant (ns). Details of the statistical analyses are presented in Supplementary Table 1.

**Supplementary Figure 4.**
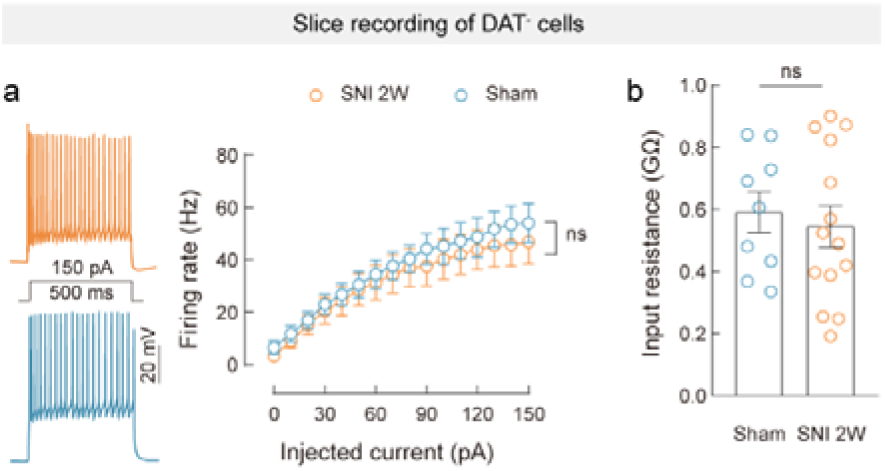
The excitability of DAT^-^ neurons in SNI 2W and Sham mice. **a**, Representative traces and statistical data of firing rate recorded from DAT^-^ neurons of SNI 2W and Sham mice. **b**, Statistical data for input resistance recorded from DAT^-^ neurons of SNI 2W and Sham mice. Significance was assessed by two-way ANOVA followed by Bonferroni’s multiple comparisons test in (**a**) and two-tailed unpaired Student’s *t*-test in (**b**). All data are presented as the mean ± s.e.m. Not significant (ns). Details of the statistical analyses are presented in Supplementary Table 1.

**Supplementary Figure 5.**
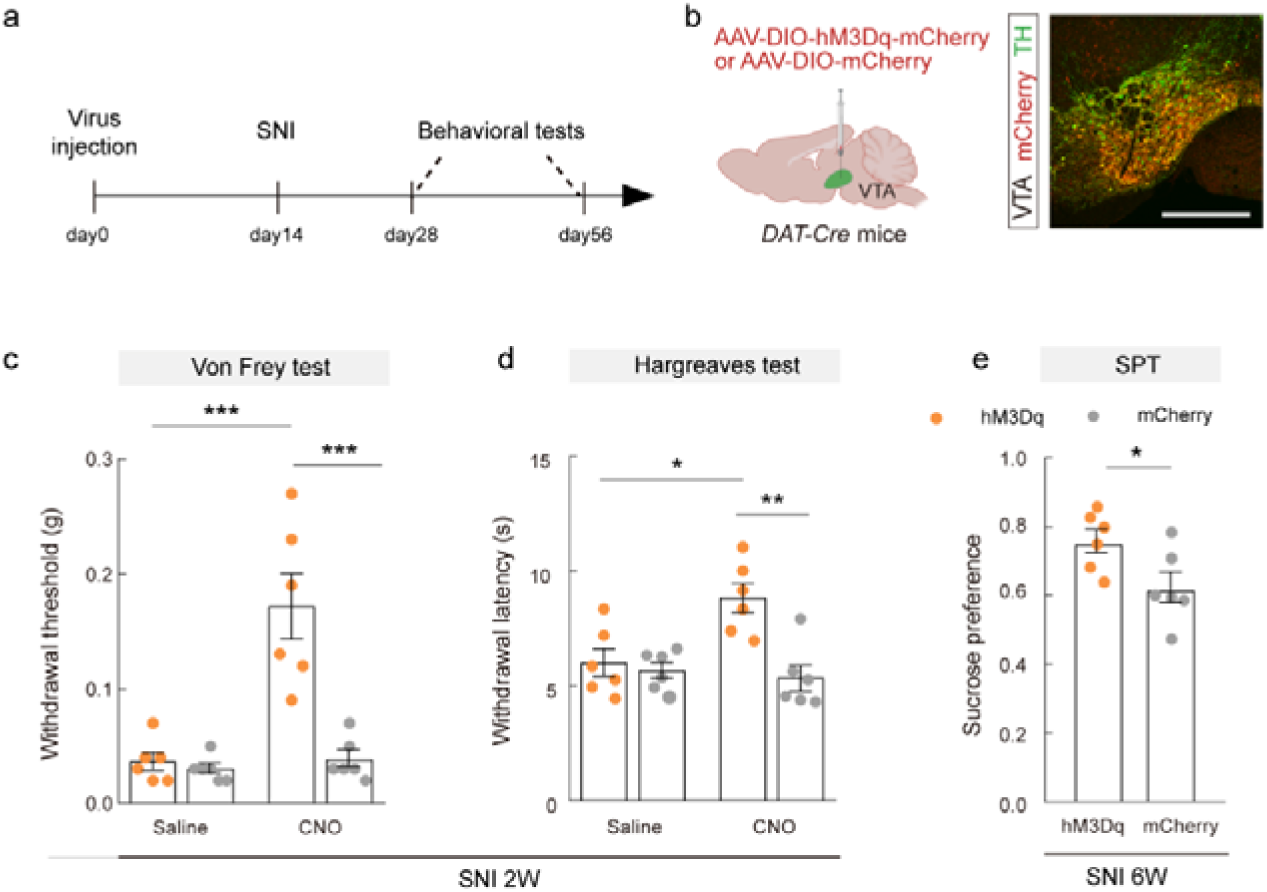
Effects of chemogenetic activation of DA^VTA^ neurons on chronic pain. **a**, Schematic of the experimental design. **b**, Schematic of VTA injection of AAV-DIO-hM3Dq-mCherry/AAV-DIO-mCherry (left) in *DAT*-Cre mice and representative images of mCherry-expressed neurons co-localized with TH immunofluorescence in the VTA (right). Scale bar, 500 μm. **c**-**e**, Statistical data for mechanical paw withdrawal threshold (**c**), thermal paw withdrawal latency (**d**) and SPT (**e**) of hM3Dq-mCherry-expressing and mCherry-expressing SNI mice with saline or CNO treatment. Significance was assessed by two-way ANOVA followed by Bonferroni’s multiple comparisons test in (**c**, **d**) and two-tailed unpaired Student’s *t*-test in (**e**). All data are presented as the mean ± s.e.m. \**P* < 0.05, \*\**P* < 0.01, \*\*\**P* < 0.001. Details of the statistical analyses are presented in Supplementary Table 1.

**Supplementary Figure 6.**
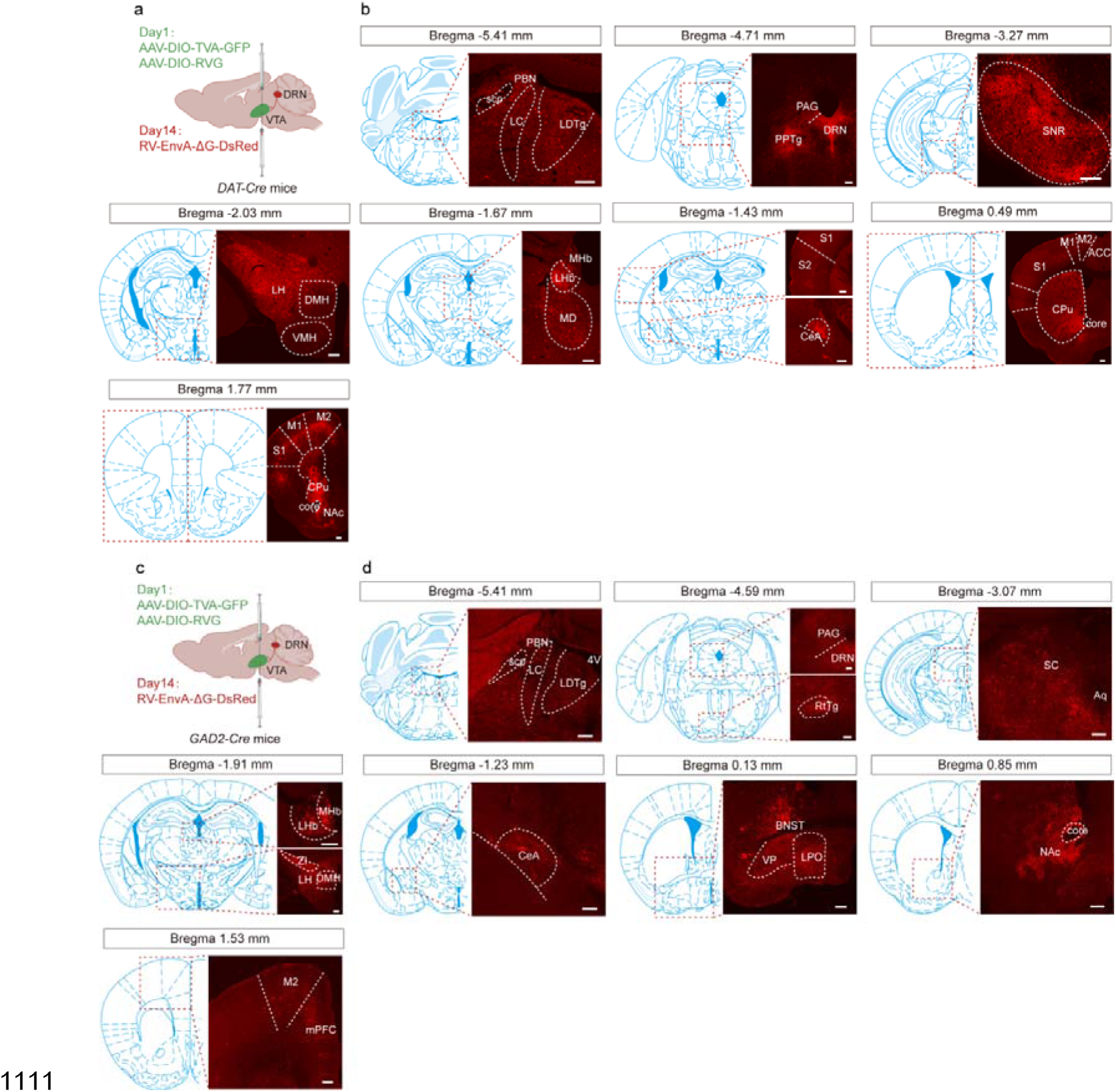
Mapping of monosynaptic inputs to DA^VTA^ and GABA^VTA^ neurons. **a**, **c**, Schematic of the Cre-dependent retrograde monosynaptic tracing strategy in *DAT*-Cre and *GAD2*-Cre mice. **b**, **d**, Sample images showing DsRed-expressing cells (red) that make monosynaptic inputs onto DA^VTA^ neurons or GABA^VTA^ neurons. Scale bars, 200 μm. PBN, parabrachial nucleus; LC, locus coeruleus; LDTg, laterodorsal tegmentum; PAG, periaqueductal gray; DRN, dorsal raphe nucleus; PPTg, pedunculopontine tegmentum; SNR, substantia nigra pars reticulata; LH, lateral hypothalamus; DMH, dorsomedial hypothalamus; VMH, ventromedial hypothalamus; LHb, lateral habenular nucleus; MHb, medial habenular nucleus; MD, mediodorsal thalamic nucleus; CeA, central nucleus of the amygdala; CPu, caudate putamen; S1, primary somatosensory cortex; S2, secondary somatosensory cortex; M1, primary motor cortex; M2, secondary motor cortex. ACC, anterior cingulate cortex; NAc, nucleus accumbens; BNST, bed nucleus of the stria terminalis; RtTg, reticulotegmental nucleus of the pons; ZI, zona incerta; SC, superior colliculus; VP, ventral pallidum; LPO, lateral preoptic area; mPFC, medial prefrontal cortex.

**Supplementary Figure 7.**
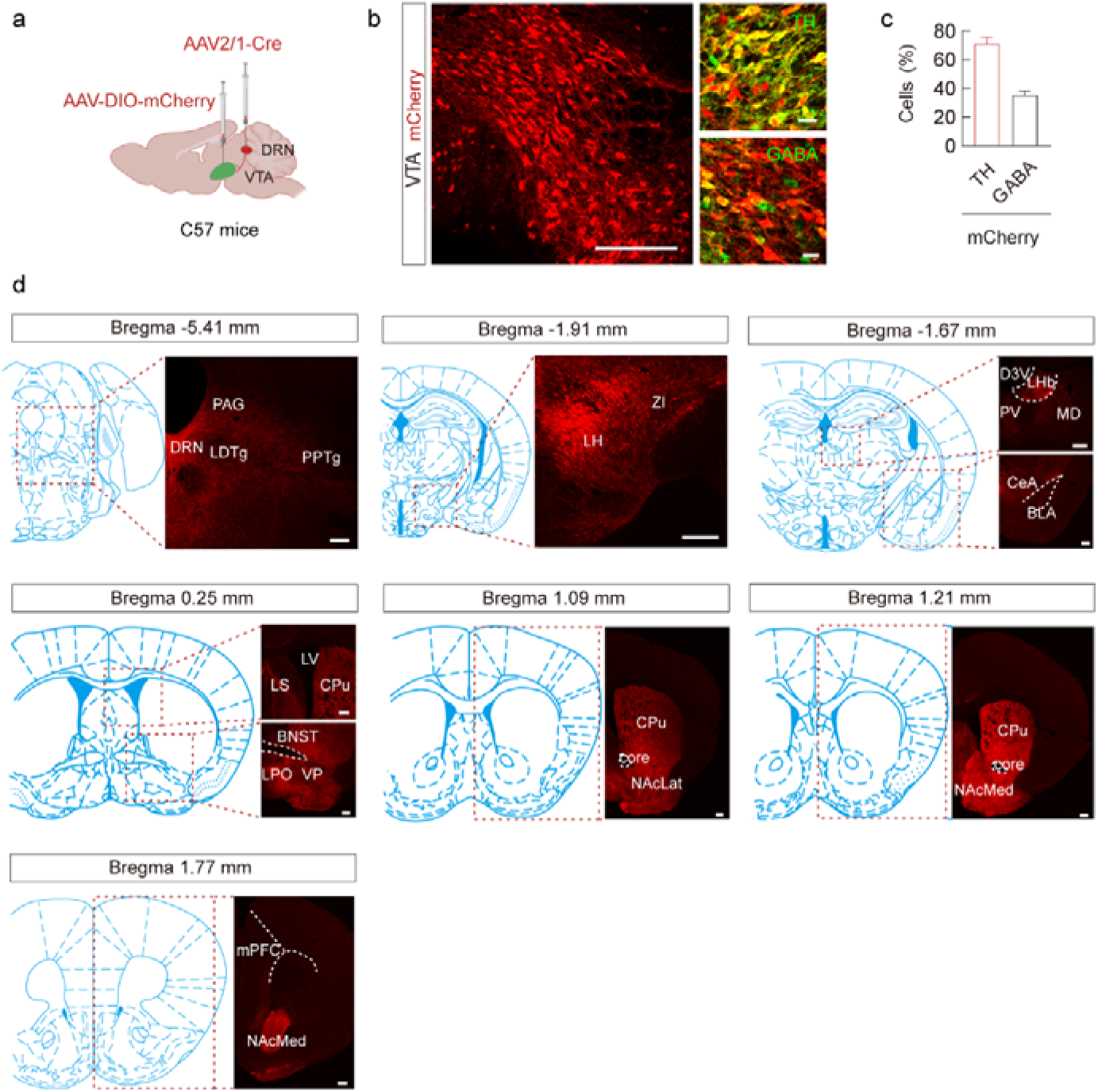
Cre-dependent anterograde trans-monosynaptic tracing of DRN neurons. **a**, Schematic of the Cre-dependent anterograde trans-monosynaptic tracing strategy in *C57* mice. **b**, Typical images showing mCherry expression within the VTA. Scale bars, 200 μm (left) and 100 μm (right). **c**, Summary data for percentage of mCherry-labeled neurons co-localized with TH and GABA immunofluorescence within the VTA, n = 6 sections from three mice (right). **d**, VTA neural terminals expressing mCherry in different brain regions. Scale bars, 200 μm. PBN, parabrachial nucleus; LDTg, laterodorsal tegmentum; PAG, periaqueductal gray; DRN, dorsal raphe nucleus; PPTg, pedunculopontine tegmentum; LH, lateral hypothalamus; ZI, zona incerta; LHb, lateral habenular nucleus; PV, paraventricular thalamic nucleus; MD, mediodorsal thalamic nucleus; BLA, basolateral amygdala; CeA, central nucleus of the amygdala; LS, lateral septal nucleus; CPu, caudate putamen; LPO, lateral preoptic area; BNST, bed nucleus of the stria terminalis; VP, ventral pallidum; NAcLat, NAc lateral shell; NAcMed, NAc medial shell; mPFC, medial prefrontal cortex.

**Supplementary Figure 8.**
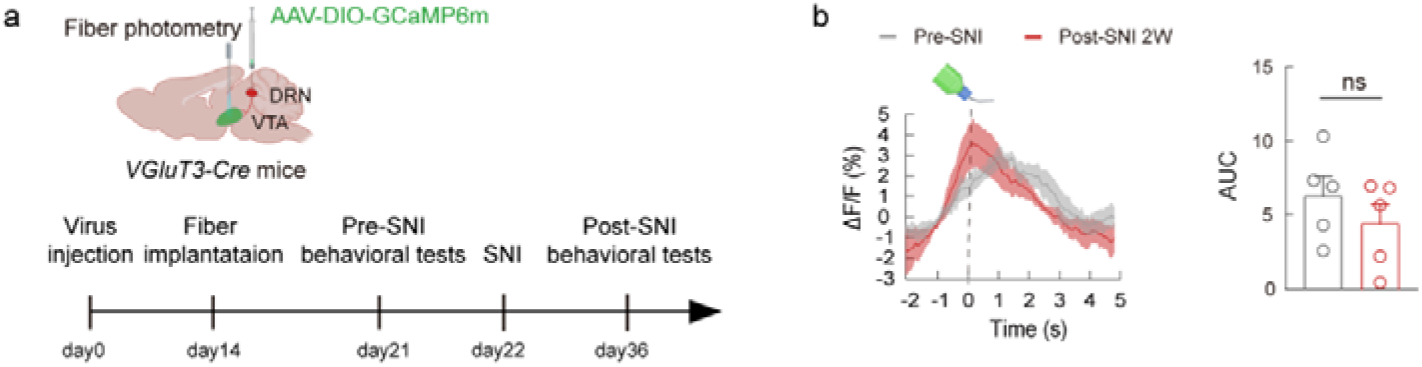
Ca^2+^ signal of VGluT3^DRN→VTA^ afferents in post-SNI 2W mice. **a**, Schematic of the experimental design and schematic diagram of fiber photometry of *VGluT3-Cre* mice. **b**, Averaged responses (left) and AUC during 0-5 s (right) showing Ca^2+^ responses evoked by sucrose licking in SNI 2W mice compared with pre-SNI mice. Significance was assessed by two-tailed paired Student’s *t*-test in (**b**). Not significant (ns). Details of the statistical analyses are presented in Supplementary Table 1.

**Supplementary Figure 9.**
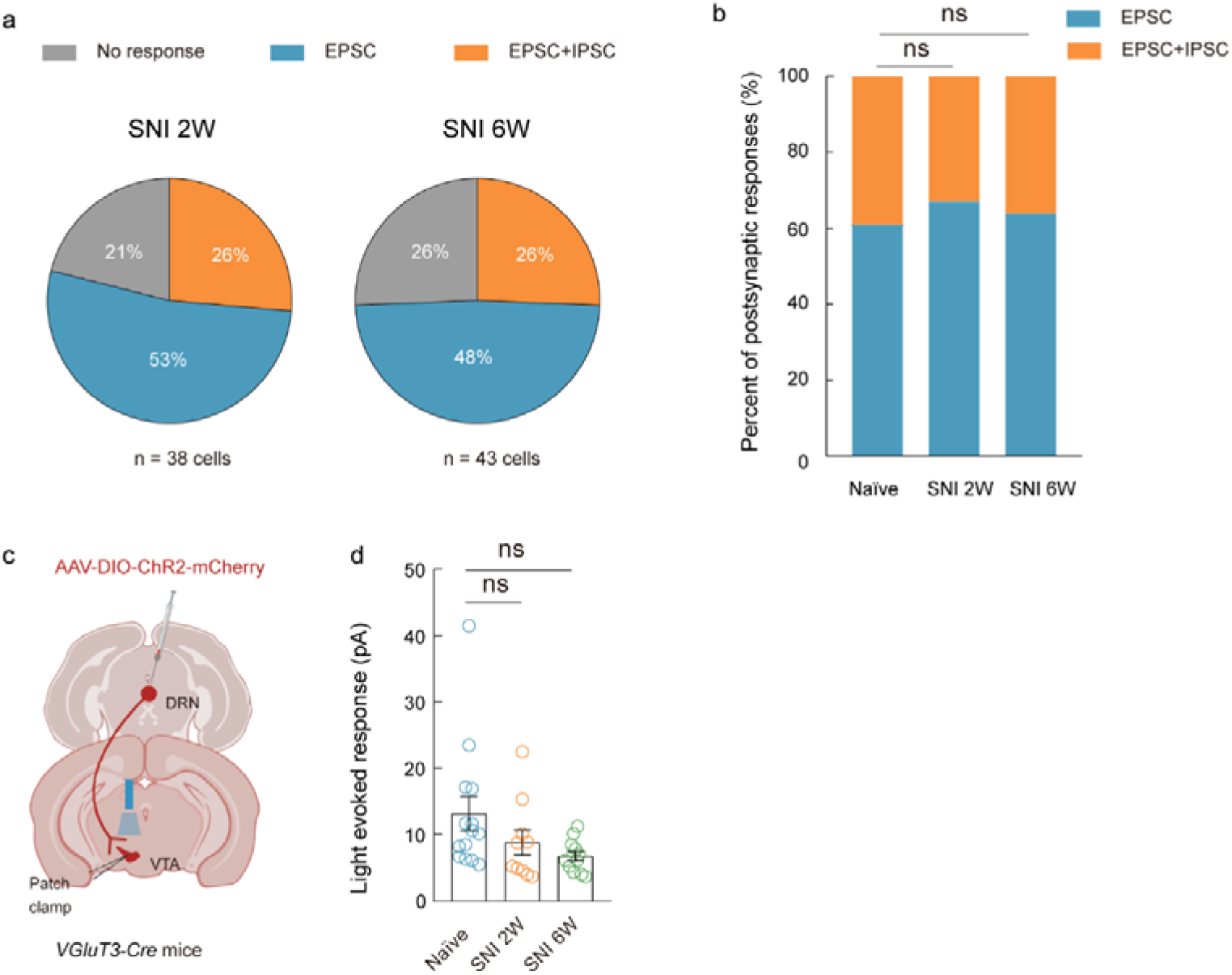
Analysis of nerve injury-induced changes in the postsynaptic responses of VGluT3^DRN^-targeted VTA. **a**, Pie chart showing the distribution of light-evoked response types in VGluT3^DRN^-targeted VTA neurons of SNI 2W and SNI 6W mice. **b**, Bar graph illustrating the percentage of EPSC- and EPSC+IPSC-type responses in all postsynaptic responses recorded from naïve, SNI 2W and SNI 6W mice. **c**, Schematic showing viral injection and the electrophysiological recordings in acute slices from *VGluT3-Cre* mice. **d,** Statistical data for the amplitudes of light-evoked slow IPSCs in VGluT3^DRN^-targeted VTA neurons of naïve, SNI 2W and SNI 6W mice. Significance was assessed by Chi-square test between groups in (**a**), Fisher’s exact test in (**b**), and one-way ANOVA followed by Bonferroni’s multiple comparisons test in (**d**). All data are presented as the mean ± s.e.m. Not significant (ns). Details of the statistical analyses are presented in Supplementary Table 1.

**Supplementary Figure 10.**
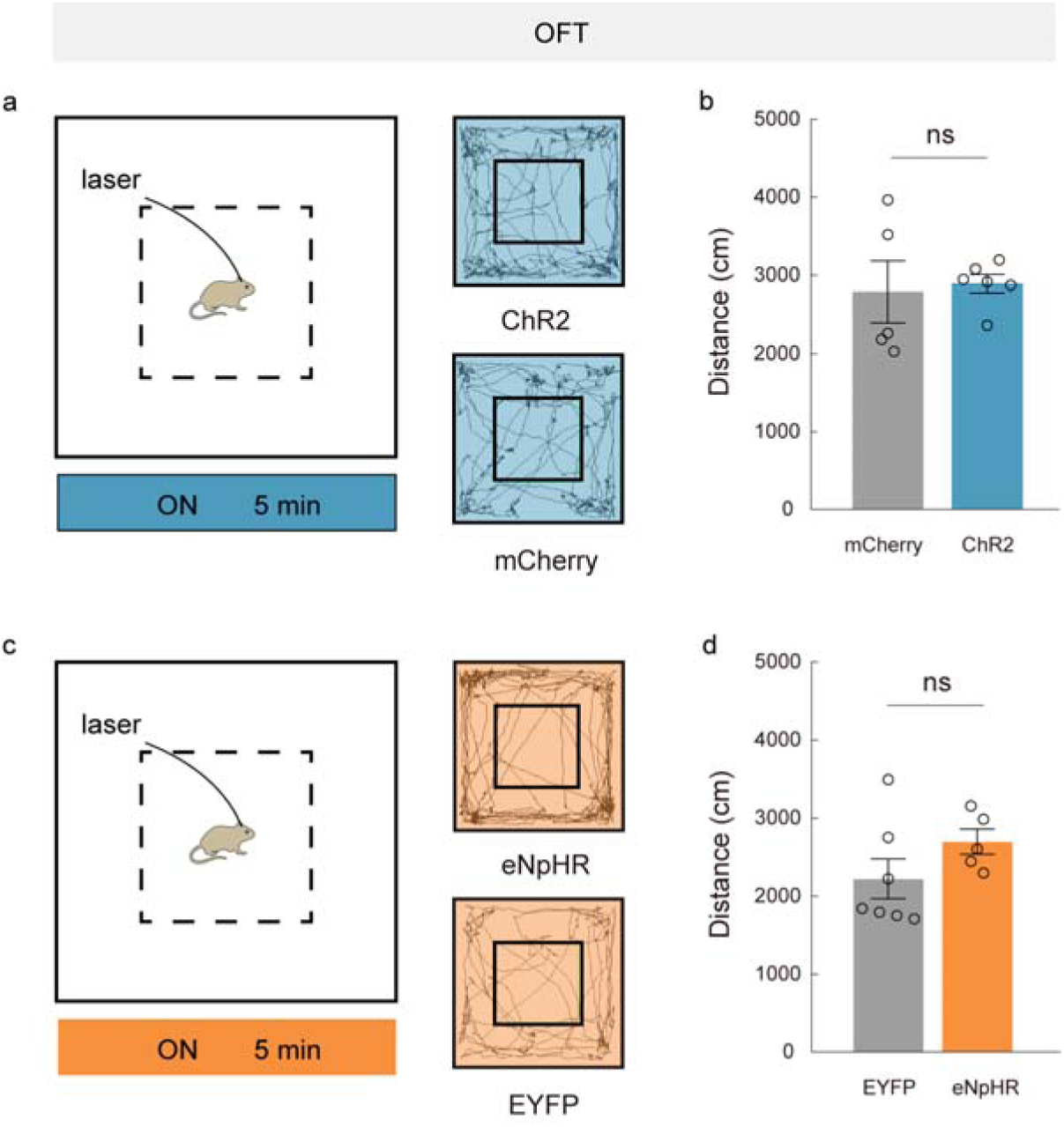
Motor activity in mice with light stimulation of VGluT3^DRN→VTA^ afferents. **a**, **c** Experimental design (left) and representative trajectories (right) of animals expressing ChR2/mCherry (**a**) or eNpHR/EYFP (**c**) in VGluT3^DRN^ neurons during VTA light stimulation. **b**, **d**, Mean distance of ChR2/mCherry-expressing (b) and eNpHR/EYFP-expressing (d) mice with 20 Hz stimulation of VGluT3^DRN→VTA^ afferents in OFT. Significance was assessed by two-tailed unpaired Student’s *t*-test in (**b**, **d**). All data are presented as the mean ± s.e.m. Not significant (ns). Details of the statistical analyses are presented in Supplementary Table 1.

**Supplementary Figure 11.**
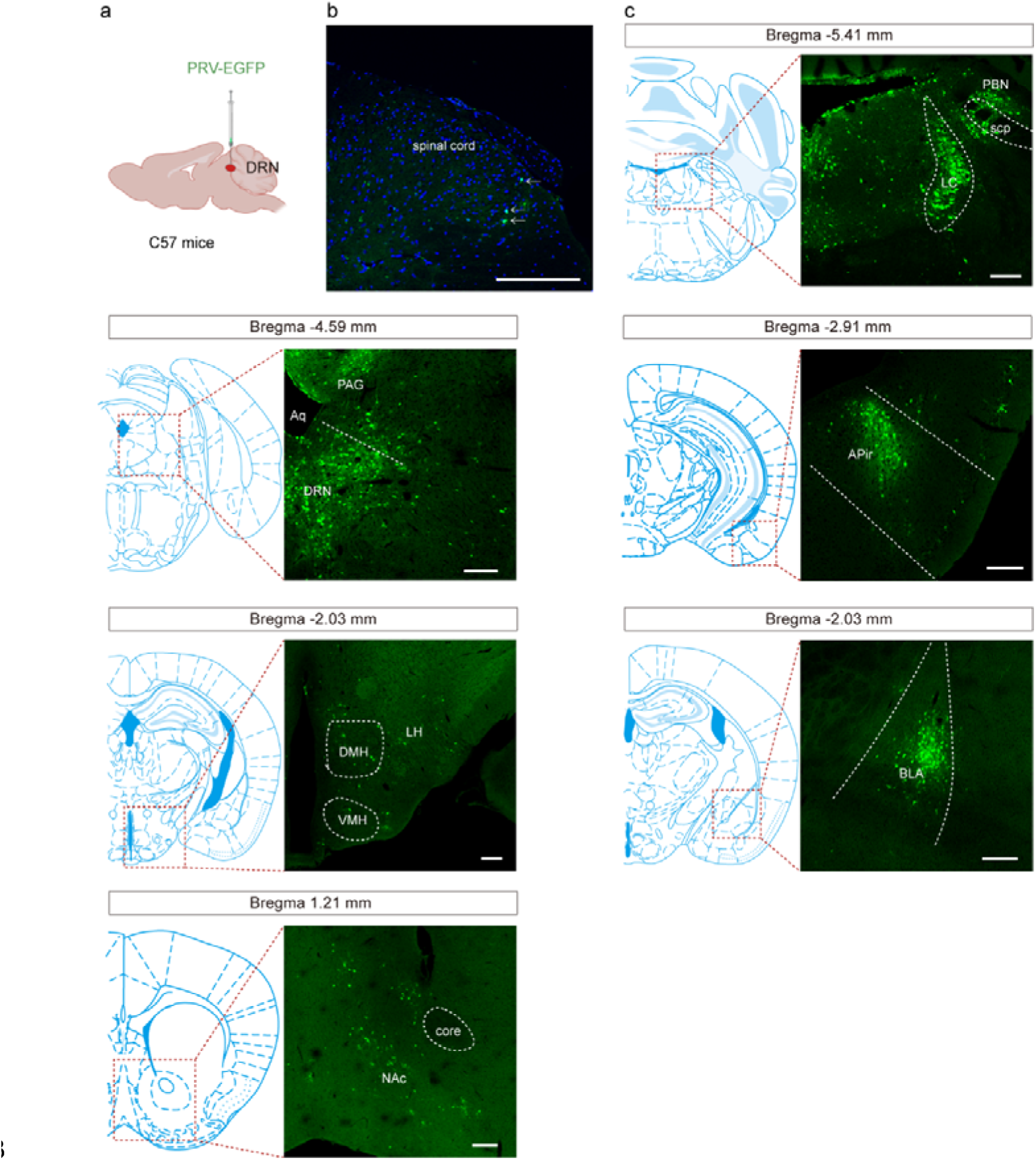
Retrograde and trans-synaptic tracing of DRN-upstream neurons with PRV-EGFP injection. **a**, Schematic of the retrograde and trans-synaptic tracing strategy using PRV-EGFP in *C57* mice. **b**, EGFP-expressing neurons detected in the dorsal horn of the spinal cord. Scale bars, 200 μm. **c**, Sample images showing EGFP-expressing cells (green) that make trans-synaptic inputs onto DRN neurons. Scale bars, 200 μm. PBN, parabrachial nucleus; PAG, periaqueductal gray; LC, locus coeruleus; DRN, dorsal raphe nucleus; APir, amygdalopiriform transition area; LH, lateral hypothalamus; DMH, dorsomedial hypothalamus; VMH, ventromedial hypothalamus; BLA, basolateral amygdala; NAc, nucleus accumbens.

**Supplementary Table 1.** Statistical analyses related to figures 1-7 and supplementary figures 1-11.

| Fig | Comparison<br>(sample size) | Analysis | *P* value | F/T/W/U<br>value |
| --- | --- | --- | --- | --- |
| Fig. 1e | Pre-SNI (6)<br>vs. Post-SNI<br>2W (6) | Paired two sample<br>t test | *P* = 0.0004 | t (5) =<br>8.392 |
| Fig. 1g | Pre-SNI (6)<br>vs. Post-SNI<br>6W (6) |  | *P* = 0.0264 | t (5) =<br>3.114 |
| Fig. 1i | Row 16 Sham<br>(23) vs. SNI<br>2W (27) | Two-way ANOVA<br>followed by<br>Bonferroni's<br>multiple<br>comparisons test | *P* = 0.0056 | F (2, 77) =<br>5.612 |
|  | Row 16 Sham<br>(23) vs. SNI<br>6W (30) |  | *P* = 0.0112 |  |
| Fig. 1j | Sham (24) vs.<br>SNI 2W (28) | One-way ANOVA<br>followed by<br>Bonferroni's<br>multiple<br>comparisons test | *P* = 0.2630 | F (2, 69) =<br>9.043 |
|  | Sham (24) vs.<br>SNI 6W (20) |  | *P* = 0.0002 |  |
| Fig. 2d | *DAT-Cre* (9)<br>vs. *GAD2-Cre*<br>(9) | Unpaired two<br>sample t test | *P* < 0.0001 | t (16) =<br>7.928 |
| Fig. 2i | ACSF (6) vs.<br>CNQX (6) | Paired two sample<br>t test | *P* = 0.0040 | t (5) =<br>5.041 |
| Fig. 2l | ACSF (5) vs.<br>Ketanserin (5) | Paired two sample<br>t test | *P* = 0.0019 | t (4) =<br>7.239 |
| Fig. 2n | DAT<sup>+</sup> (17) vs.<br>DAT<sup>-</sup> (6) | Unpaired two<br>sample t test | *P* = 0.4247 | t (21) =<br>0.8142 |
| Fig. 3d | Pre-SNI (6)<br>vs. Post-SNI<br>2W (6) | Paired two sample<br>t test | *P* = 0.0096 | t (5) =<br>4.072 |
| Fig. 3e | Pre-SNI (6)<br>vs. Post-SNI<br>6W (6) |  | *P* = 0.0072 | t (5) =<br>4.367 |
| Fig. 3h | Sham (14) vs. SNI 2W (13) | One-way ANOVA followed by Bonferroni's multiple comparisons test | $P = 0.0305$ | $F(2, 41) = 4.014$ |
| | Sham (14) vs. SNI 6W (17) | | $P = 0.0429$ | |
| Fig. 3i | Sham (20) vs. SNI 2W (18) | One-way ANOVA followed by Bonferroni's multiple comparisons test | $P = 0.0389$ | $F(2, 52) = 4.402$ |
| | Sham (20) vs. SNI 6W (17) | | $P = 0.0210$ | |
| Fig. 3k | Sham (32) vs. SNI 2W (23) | One-way ANOVA followed by Bonferroni's multiple comparisons test | $P = 0.0005$ | $F(2, 71) = 7.833$ |
| | Sham (32) vs. SNI 6W (19) | | $P = 0.0387$ | |
| Fig. 3l | Sham (32) vs. SNI 2W (23) | One-way ANOVA followed by Bonferroni's multiple comparisons test | $P = 0.0105$ | $F(2, 71) = 7.417$ |
| | Sham (32) vs. SNI 6W (19) | | $P = 0.0017$ | |
| Fig. 3m | Sham (32) vs. SNI 2W (23) | One-way ANOVA followed by Bonferroni's multiple comparisons test | $P = 0.7065$ | $F(2, 71) = 2.076$ |
| | Sham (32) vs. SNI 6W (19) | | $P = 0.3992$ | |
| Fig. 3n | Row 11 Sham (23) vs. SNI 2W (23) | Two-way ANOVA followed by Bonferroni's multiple comparisons test | $P = 0.0167$ | $F(2, 61) = 10.80$ |
| | Row 11 Sham (23) vs. SNI 6W (18) | | $P = 0.0173$ | |
| Fig. 3o | Sham (25) vs. SNI 2W (22) | One-way ANOVA followed by Bonferroni's multiple comparisons test | $P = 0.3780$ | $F(2, 62) = 4.570$ |
| | Sham (25) vs. SNI 6W (18) | | $P = 0.0071$ | |
| Fig. 3p | Sham (18) vs. SNI 2W (17) | One-way ANOVA followed by | $P = 0.0042$ | $F(2, 46) = 5.743$ |
| | Sham (18) vs.<br>SNI 6W (14) | Bonferroni's<br>multiple<br>comparisons test | $P = 0.0439$ | |
| Fig. 3q | Sham (15) vs.<br>SNI 2W (15) | One-way ANOVA<br>followed by<br>Bonferroni's<br>multiple<br>comparisons test | $P > 0.9999$ | F (2, 42) =<br>5.513 |
| | Sham (15) vs.<br>SNI 6W (15) | | $P = 0.0269$ | |
| Fig. 4c | Sham: ChR2<br>(6) OFF vs.<br>mCherry (9)<br>OFF | Two-way ANOVA<br>followed by<br>Bonferroni's<br>multiple<br>comparisons test | $P > 0.9999$ | F (1, 26) =<br>2.821 |
| | Sham: ChR2<br>(6) ON vs.<br>mCherry (9)<br>ON | | $P = 0.5255$ | |
| | Sham: ChR2<br>(6) ON vs.<br>ChR2 (6)<br>OFF | | $P > 0.9999$ | |
| | Sham:<br>mCherry (9)<br>ON vs.<br>mCherry (9)<br>OFF | | $P > 0.9999$ | |
| | SNI: ChR2<br>(9) OFF vs.<br>mCherry (6)<br>OFF | Two-way ANOVA<br>followed by<br>Bonferroni's<br>multiple<br>comparisons test | $P > 0.9999$ | F (1, 26) =<br>21.10 |
| | SNI: ChR2<br>(9) ON vs.<br>mCherry (6)<br>ON | | $P < 0.0001$ | |
| | SNI: ChR2<br>(9) ON vs.<br>ChR2 (9)<br>OFF | | $P < 0.0001$ | |
| | SNI: mCherry<br>(6) ON vs.<br>mCherry (6)<br>OFF | | $P > 0.9999$ | |
| Fig. 4d | Sham: ChR2<br>(6) OFF vs.<br>mCherry (9)<br>OFF | Two-way ANOVA<br>followed by<br>Bonferroni's<br>multiple<br>comparisons test | $P > 0.9999$ | F (1, 26) =<br>0.2520 |
| | Sham: ChR2<br>(6) ON vs.<br>mCherry (9)<br>ON | | $P > 0.9999$ | |
| | Sham: ChR2<br>(6) ON vs.<br>ChR2 (6)<br>OFF | | $P > 0.9999$ | |
| | Sham:<br>mCherry (9)<br>ON vs.<br>mCherry (9)<br>OFF | | $P > 0.9999$ | |
| | SNI: ChR2<br>(9) OFF vs.<br>mCherry (6)<br>OFF | Two-way ANOVA<br>followed by<br>Bonferroni's<br>multiple<br>comparisons test | $P > 0.9999$ | F (1, 26) =<br>2.211 |
| | SNI: ChR2<br>(9) ON vs.<br>mCherry (6)<br>ON | | $P = 0.0153$ | |
| | SNI: ChR2<br>(9) ON vs.<br>ChR2 (9)<br>OFF | | $P = 0.0005$ | |
| | SNI: mCherry<br>(6) ON vs.<br>mCherry (6)<br>OFF | | $P > 0.9999$ | |
| Fig. 4e | Sham ChR2 :<br>Pre (6) vs.<br>Test (6) | Paired two sample<br>t test | $P = 0.0470$ | $t(5) = 2.622$ |
| | Sham<br>mCherry : Pre<br>(6) vs. Test<br>(6) | | $P = 0.7192$ | $t(5) = 0.3806$ |
| | SNI ChR2 :<br>Pre (10) vs.<br>Test (10) | | $P = 0.0118$ | $t(9) = 3.146$ |
| | SNI<br>mCherry : Pre<br>(5) vs. Test<br>(5) | | $P = 0.4282$ | $t(4) = 0.8807$ |
| Fig. 4f | ChR2 (6)<br>Sham vs.<br>mCherry (6)<br>Sham | Two-way ANOVA<br>followed by<br>Bonferroni's<br>multiple<br>comparisons test | $P = 0.7663$ | $F(1, 20) = 2.215$ |
| | ChR2 (7) SNI<br>vs. mCherry<br>(5) SNI | | $P = 0.0094$ | |
| | ChR2 (7) SNI<br>vs. ChR2 (6)<br>Sham | | $P > 0.9999$ | |
| | mCherry (5)<br>SNI vs.<br>mCherry (6)<br>Sham | | $P = 0.0047$ | |
| Fig. 4i | ACSF:<br>hM3Dq (6)<br>vs. mCherry<br>(5) | Two-way ANOVA<br>followed by<br>Bonferroni's<br>multiple<br>comparisons test | $P < 0.0001$ | $F(1, 27) = 150.6$ |
| | CNQX:<br>hM3Dq (6)<br>vs. mCherry | | $P > 0.9999$ | |
|  | (5) |  |  |  |
| | Ketanserin:<br>hM3Dq (6)<br>vs. mCherry<br>(5) | | $P < 0.0001$ | |
| | hM3Dq:<br>ACSF (6) vs.<br>CNQX (6) | | $P < 0.0001$ | |
| | hM3Dq:<br>ACSF (6) vs.<br>Ketanserin (6) | | $P > 0.9999$ | |
| | hM3Dq:<br>CNQX (6) vs.<br>Ketanserin (6) | | $P < 0.0001$ | |
| Fig. 4j | ACSF:<br>hM3Dq (7)<br>vs. mCherry<br>(6) | Two-way ANOVA<br>followed by<br>Bonferroni's<br>multiple<br>comparisons test | $P = 0.0035$ | F (1, 33) =<br>19.59 |
| | CNQX:<br>hM3Dq (7)<br>vs. mCherry<br>(6) | | $P > 0.9999$ | |
| | Ketanserin:<br>hM3Dq (7)<br>vs. mCherry<br>(6) | | $P = 0.0473$ | |
| | hM3Dq:<br>ACSF (7) vs.<br>CNQX (7) | | $P = 0.0259$ | |
| | hM3Dq:<br>ACSF (7) vs.<br>Ketanserin (7) | | $P > 0.9999$ | |
| | hM3Dq:<br>CNQX (7) vs.<br>Ketanserin (7) | | $P = 0.0058$ | |
| Fig. 5d | OFF: eNpHR<br>(9) vs.<br>EYFP (8) | Two-way ANOVA<br>followed by<br>Bonferroni's<br>multiple<br>comparisons test | $P = 0.4427$ | $F(1, 30) = 87.33$ |
| | eNpHR: OFF<br>(9) vs. ON (9) | | $P < 0.0001$ | |
| | EYFP: OFF<br>(8) vs. ON (8) | | $P > 0.9999$ | |
| | ON: eNpHR<br>(9) vs. EYFP<br>(8) | | $P < 0.0001$ | |
| Fig. 5e | OFF: eNpHR<br>(11) vs.<br>EYFP (8) | Two-way ANOVA<br>followed by<br>Bonferroni's<br>multiple<br>comparisons test | $P > 0.9999$ | $F(1, 34) = 24.74$ |
| | eNpHR: OFF<br>(11) vs. ON<br>(9) | | $P < 0.0001$ | |
| | EYFP: OFF<br>(8) vs. ON (8) | | $P = 0.3993$ | |
| | ON: eNpHR<br>(9) vs. EYFP<br>(8) | | $P < 0.0001$ | |
| Fig. 5f | eNpHR : Pre<br>(8) vs. Test<br>(8) | Paired two sample<br>t test | $P = 0.0003$ | $t(7) = 6.576$ |
| | EYFP :Pre (8)<br>vs. Test (8) | | $P = 0.6639$ | $t(7) = 0.4535$ |
| Fig. 5i | hM4Di (5) vs<br>mCherry (5) | Two-way ANOVA<br>followed by<br>Bonferroni's<br>multiple<br>comparisons test | | $F(1, 8) = 704.4$ |
| | BL | | $P > 0.9999$ | |
| | 21 | | $P > 0.9999$ | |
| | 23 | | $P = 0.0045$ | |
| | 25 | | $P = 0.0008$ | |
| | 27 | | $P = 0.0096$ | |
| | 29 | | $P = 0.0168$ | |
| | 35 | | $P = 0.0041$ | |
| | 42 | | $P = 0.0121$ | |
| | 49 | | $P = 0.0005$ | |
| | 56 | | $P = 0.0147$ | |
| | 63 | | $P = 0.0012$ | |
| Fig. 5j | hM4Di : Pre<br>(5) vs. Test<br>(5) | Paired two sample<br>t test | $P = 0.0112$ | t (4) =<br>4.461 |
| | mCherry : Pre<br>(5) vs. Test<br>(5) | | $P = 0.7386$ | t (4) =<br>0.3578 |
| Fig. 5k | hM3Di (5) vs.<br>mCherry (5) | Unpaired two<br>sample t test | $P = 0.0442$ | t (8) =<br>2.386 |
| Fig. 6c | Pre-SNI (6)<br>vs. Post-SNI<br>2W (6) | Paired two sample<br>t test | $P = 0.0027$ | t (5) =<br>5.507 |
| Fig. 6d | Pre-SNI (5)<br>vs. Post-SNI<br>2W (5) | | $P = 0.7926$ | t (4) =<br>0.2811 |
| Fig. 6e | Pre-SNI (6)<br>vs. Post-SNI<br>2W (6) | | $P = 0.0001$ | t (5) =<br>10.58 |
| Fig. 6f | Pre-SNI (5)<br>vs. Post-SNI<br>2W (5) | | $P = 0.3838$ | t (4) =<br>0.9773 |
| Fig. 6g | Pre-SNI (6)<br>vs. Post-SNI | | $P = 0.0202$ | t (5) =<br>3.358 |
|  | 6W (6) |  |  |  |
| Fig. 6h | Pre-SNI (5)<br>vs. Post-SNI<br>6W (5) | | $P = 0.9069$ | $t(4) = 0.1246$ |
| Fig. 7c | ACSF: OFF (5) vs. ON (5) | Two-way ANOVA followed by Bonferroni's multiple comparisons test | $P < 0.0001$ | $F(1, 24) = 66.98$ |
| | SCH23390: OFF (5) vs. ON (5) | | $P < 0.0001$ | |
| | Eticlopride: OFF (5) vs. ON (5) | | $P > 0.9999$ | |
| | ON: ACSF (5) vs. SCH23390 (5) | | $P > 0.9999$ | |
| | ON: ACSF (5) vs. Eticlopride (5) | | $P < 0.0001$ | |
| | ON: SCH23390 (5) vs. Eticlopride (5) | | $P < 0.0001$ | |
| Fig. 7d | ACSF (5) vs. SCH23390 (5) | One-way ANOVA followed by Bonferroni's multiple comparisons test | $P = 0.037$ | $F(2, 12) = 5.633$ |
| | ACSF (5) vs. Eticlopride (5) | | $P = 0.8775$ | |
| Fig. 7e | ACSF: OFF (6) vs. ON (6) | Two-way ANOVA followed by Bonferroni's multiple comparisons test | $P < 0.0001$ | $F(1, 30) = 236.9$ |
| | SCH23390: OFF (6) vs. ON (6) | | $P < 0.0001$ | |
| | Eticlopride:<br>OFF (6) vs.<br>ON (6) | | $P < 0.0001$ | |
| | ON: ACSF<br>(6) vs.<br>SCH23390<br>(6) | | $P > 0.9999$ | |
| | ON: ACSF<br>(6) vs.<br>Eticlopride<br>(6) | | $P > 0.9999$ | |
| | ON:<br>SCH23390<br>(6) vs.<br>Eticlopride<br>(6) | | $P > 0.9999$ | |
| Fig. 7f | ACSF (6) vs.<br>SCH23390<br>(6) | One-way ANOVA<br>followed by<br>Bonferroni's<br>multiple<br>comparisons test | $P = 0.9983$ | F (2, 15) =<br>0.1188 |
| | ACSF (6) vs.<br>Eticlopride<br>(6) | | $P = 0.8684$ | |
| Supplementary<br>Fig. 1b |  |  |  |  |
|  | Sham (6) vs<br>SNI (5) | Two-way RM<br>ANOVA with<br>Bonferroni post<br>hoc analysis |  | F (1, 9) =<br>378.6 |
| | 0d | | $P > 0.9999$ | |
| | 7d | | $P = 0.0009$ | |
| | 14d | | $P = 0.0003$ | |
| | 21d | | $P = 0.0005$ | |
| | 27d | | $P = 0.0002$ | |
| | 35d | | $P = 0.0034$ | |
| | 42d | | $P = 0.0009$ | |
| Supplementary<br>Fig. 1c | 2W Sham (6)<br>vs SNI (6) | Unpaired two<br>sample t test | $P = 0.3892$ | t (10) =<br>0.9001 |
| | 6W Sham (6)<br>vs SNI (6) | | $P = 0.0312$ | t (10) =<br>2.504 |
| Supplementary<br>Fig. 2a | Pre-SNI (6)<br>vs. Post-SNI<br>(6) | Two-way RM<br>ANOVA with<br>Bonferroni post<br>hoc analysis |  |  |
| | 0.02 | | $P = 0.0007$ | F (1, 10) =<br>453.8 |
| | 0.04 | | $P = 0.001$ | |
| | 0.07 | | $P < 0.0001$ | |
| | 0.16 | | $P < 0.0001$ | |
| | 0.4 | | $P < 0.0001$ | |
| | 0.6 | | $P = 0.0001$ | |
| | 1 | | $P = 0.1538$ | |
| Supplementary<br>Fig. 2b | Pre-SNI : Pre<br>(6) vs. Test<br>(6) | Paired two sample<br>t test | $P = 0.0995$ | t (5) =<br>2.019 |
| | Post-SNI : Pre<br>(6) vs. Test<br>(6) | | $P = 0.0237$ | t (5) =<br>3.209 |
| Supplementary<br>Fig. 3b | Pre-SNI (3)<br>vs. Post-SNI | Paired two sample<br>t test | $P = 0.2400$ | t (2) =<br>1.654 |
|  | 2W (3) |  |  |  |
| Supplementary<br>Fig. 4a | Row 16 Sham<br>(8) vs. SNI<br>2W (11) | Two-way ANOVA<br>followed by<br>Bonferroni's<br>multiple<br>comparisons test | $P > 0.9999$ | F (1, 17) =<br>0.2474 |
| Supplementary<br>Fig. 4b | Sham (9) vs.<br>SNI 2W (14) | Unpaired two<br>sample t test | $P = 0.6418$ | t (21) =<br>0.4719 |
| Supplementary<br>Fig. 5c | Saline:<br>hM3Dq (6)<br>vs. mCherry<br>(6) | Two-way ANOVA<br>followed by<br>Bonferroni's<br>multiple<br>comparisons test | $P > 0.9999$ | F (1, 20) =<br>20.63 |
| | CNO: hM3Dq<br>(6) vs.<br>mCherry (6) | | $P < 0.0001$ | |
| | hM3Dq:<br>Saline (6) vs.<br>CNO (6) | | $P < 0.0001$ | |
| | mCherry:<br>Saline (6) vs.<br>CNO (6) | | $P > 0.9999$ | |
| Supplementary<br>Fig. 5d | Saline:<br>hM3Dq (6)<br>vs. mCherry<br>(6) | Two-way ANOVA<br>followed by<br>Bonferroni's<br>multiple<br>comparisons test | $P > 0.9999$ | F (1, 20) =<br>12.13 |
| | CNO: hM3Dq<br>(6) vs.<br>mCherry (6) | | $P = 0.0014$ | |
| | hM3Dq:<br>Saline (6) vs.<br>CNO (6) | | $P = 0.0102$ | |
| | mCherry:<br>Saline (6) vs.<br>CNO (6) | | $P > 0.9999$ | |
| Supplementary<br>Fig. 5e | hM3Dq (6)<br>vs. mCherry | Unpaired two<br>sample t test | $P = 0.0374$ | t (10) =<br>2.399 |
|  | (6) |  |  |  |
| Supplementary<br>Fig. 8b | Pre-SNI (5)<br>vs. Post-SNI<br>2W (5) | Paired two sample<br>t test | $P = 0.2607$ | $t(8) = 1.309$ |
| Supplementary<br>Fig. 9a | naïve vs. SNI<br>2W | Chi-square test | $P = 0.8502$ | $\chi^2 = 0.3246, df = 2$ |
| | naïve vs. SNI<br>6W | | $P = 0.6552$ | $\chi^2 = 0.8456, df = 2$ |
| Supplementary<br>Fig. 9b | naïve vs. SNI<br>2W | Fisher's exact test | $P = 0.4615$ | |
| | naïve vs. SNI<br>6W | | $P = 0.7703$ | |
| Supplementary<br>Fig. 9d | Sham (14) vs.<br>SNI 2W (10) | One-way ANOVA<br>followed by<br>Bonferroni's<br>multiple<br>comparisons test | $P = 0.2866$ | $F(2, 32) = 2.737$ |
| | Sham (14) vs.<br>SNI 6W (11) | | $P = 0.0609$ | |
| Supplementary<br>Fig. 10b | mCherry (5)<br>vs. Chr2 (6) | Unpaired two<br>sample t test | $P = 0.7904$ | $t(9) = 0.2738$ |
| Supplementary<br>Fig. 10d | EYFP (7) vs.<br>eNpHR (5) | Unpaired two<br>sample t test | $P = 0.1855$ | $t(10) = 1.422$ |

